# Characterizing and explaining impact of disease-associated mutations in proteins without known structures or structural homologues

**DOI:** 10.1101/2021.11.17.468998

**Authors:** Neeladri Sen, Ivan Anishchenko, Nicola Bordin, Ian Sillitoe, Sameer Velankar, David Baker, Christine Orengo

## Abstract

Mutations in human proteins lead to diseases. The structure of these proteins can help understand the mechanism of such diseases and develop therapeutics against them. With improved deep learning techniques such as RoseTTAFold and AlphaFold, we can predict the structure of proteins even in the absence of structural homologues. We modeled and extracted the domains from 553 disease-associated human proteins without known protein structures or close homologues in the Protein Databank (PDB). We noticed that the model quality was higher and the RMSD lower between AlphaFold and RoseTTAFold models for domains that could be assigned to CATH families as compared to those which could only be assigned to Pfam families of unknown structure or could not be assigned to either. We predicted ligand-binding sites, protein-protein interfaces, conserved residues in these predicted structures. We then explored whether the disease-associated missense mutations were in the proximity of these predicted functional sites, if they destabilized the protein structure based on ddG calculations or if they were predicted to be pathogenic. We could explain 80% of these disease-associated mutations based on proximity to functional sites, structural destabilization or pathogenicity. When compared to polymorphisms a larger percentage of disease associated missense mutations were buried, closer to predicted functional sites, predicted as destabilising and/or pathogenic. Usage of models from the two state-of-the-art techniques provide better confidence in our predictions, and we explain 93 additional mutations based on RoseTTAFold models which could not be explained based solely on AlphaFold models.

## Introduction

Mutations in genomes can either be in the coding regions or non-coding regions. A missense mutation (change in the amino acid to another) in the coding region of the genome can lead to changes in/near catalytic site residues [1,2], ligand binding sites [3,4], protein-protein interface sites [5,6], allosteric sites [7] etc; modifying the protein or rendering it inactive. The sequence of amino acids dictates the structure of the protein [8]. Some missense mutations can hence lead to changes in the structure of the protein [9,10], which make the protein non-functional. Mutations can also be found in noncoding regions of the genome. These largely affect protein production by initiating it at the wrong time and/or wrong place or reduce/eliminate the production of proteins. These lead to various diseases such as congenital heart diseases, pancreatic agenesis, campomelic dysplasia, developmental disorders and are very commonly found in cancer [11–14].

Single nucleotide polymorphisms (SNPs) are common in genes, some of which lead to disease and are being collated in databases such as dbSNP [15] and the 1000 genomes project [16]. Databases such as ClinVar [17] link mutations in humans with phenotypes. Specific databases such as COSMIC [18], OncoVar [19], DoCM [20] contain information on cancer related mutations. Several other specialized databases containing subsets of these mutations exist, such as KinMutBase (for human disease-associated protein kinase mutations) [21], ActiveDriverDB (mutations at protein post-translational modification sites) [22] among others. UniProt-KB [23] and PDBe-KB [24] also contain experimental and computational annotations of mutations and associated phenotypes. Humsavar [23] is a database of human variants from literature reports and therefore tends to have more substantial annotations from experimental evidence. DBSAV [25] is a database that contains all SNPs in the human proteome and a predicted score of their deleteriousness.

Protein structures can help gain insights into the mechanism of these disease-associated mutations. However experimental determination of protein structures is time-consuming, expensive and difficult. With the increase in the size of proteins and complexes, it becomes increasingly challenging to experimentally determine their structures. Hence, computational models can help in such cases. Homology modelling techniques (such as MODELLER [26,27], SWISS-MODEL [28] etc) can be used to model sequences in presence of structural homologues having sequence identity greater than or equal to 30%, but a large number of protein sequences have no close homologues. *Ab initio* protein structure modelling techniques (ROSETTA [29], I-TASSER [30] etc) are used in the absence of structural homologues. More recently deep neural networks trained on the massive body of available protein sequence and structure data have been shown to significantly outperform traditional homology modeling or *ab initio* approaches in terms of the accuracy of the resulting model (e.g. RaptorX [31], DMPFold [32], trRosetta [33], AlphaFold.v1 [34] etc). Models like AlphaFold.v2 [35] and RoseTTAFold [36] trained end-to-end to directly predict 3D model coordinates were shown to be able to reach near experimental accuracy. AlphaFold was shown to be the best performing technique for protein structure prediction in CASP14 (https://predictioncenter.org/casp14/). In July 2021, AlphaFold released the structures of the entire reference proteome of 21 organisms [37], which had about 50% of the residues (which could not be modeled using homology) predicted with confidence [38]. Recent studies have shown that these high-quality models can be used to predict binding sites and effects of mutations and these calculations/predictions are similar to those obtained from experiments/experimental structures [38].

Along with the protein structure, experimental and computational determination of functional residues can help explain the effect of mutations on proteins. Mutations on/near functional sites are likely to affect the functioning of the proteins. The current strategies for the computational prediction of functional sites such as catalytic sites, ligand binding sites, allosteric sites, protein-protein interaction sites etc have been reviewed elsewhere [39–42]. In addition to the predictions of functional sites, the energetic effect of mutations can also be calculated using various tools [43–46], that determine the free energy change of the mutation. These tools can predict if the mutation will destabilise the protein structure, possibly affecting its functioning. The likely deleteriousness of these mutations can be calculated based on various machine learning based pathogenicity predictors such as CADD, EVE, MutPred2 etc [47–49].

Homology models have been previously used to explain disease-associated mutations [50,51]. In this manuscript, we model and exploit RoseTTAFold and AlphaFold models of human proteins without known protein structures or homologues in the PDB [52] (sequence identity<30%) and explain the effect of deleterious missense mutations by checking if these mutations are on/near a predicted functional site, conserved site, lead to protein structure destabilization or are pathogenic.

## Results

### 1. Protein domains for disease-associated proteins

For our analysis, we considered 553 human disease-associated proteins without known protein structures or close homologues in the PDB (see Methods section 1a for criteria used to identify close homologues). These proteins were selected from the VarSite [53] database, containing disease site information from Uniprot, ClinVar and gnomAD (See Methods Section 1a for further details on protein selection). We checked the PANTHER [54–56] protein classes these proteins fall into. Of the 553 proteins, only 243 proteins have been associated with 20 out of 24 PANTHER classes. The most highly represented protein classes are metabolite interconverting proteins, transporters, and scaffold/adaptor proteins, containing 71, 36 and 16 proteins respectively (Supplementary Table2). These are followed by chaperons, protein modifying enzymes and membrane traffic proteins containing 13 proteins each. We performed gene ontology analysis using PANTHER, which showed enrichment of terms that can be associated with transport or metabolite interconverting enzymes (Details about gene ontology in Supplementary Text1). However, we would like to point out that the dataset is somewhat biased in the sense that these proteins are those without any homologues in the PDB. It is possible that many of them are therefore difficult to crystallise.

Out of the 553 proteins, 198 proteins mapped to CATH, of the remaining, 341 mapped to Pfam leaving 14 proteins that could not be assigned to CATH or Pfam domains. CATH [57,58] and Pfam [59] superfamilies can be functionally diverse, with 62% of the protein space being covered by the largest 200 CATH superfamilies [60]. Hence, only some of the members of these superfamilies will function similarly, requiring the need to subclassify them into functionally similar families called FunFams [61] (Supplementary Text13 for details). The 198 proteins assigned to CATH comprised 309 domains which were assigned to 297 FunFams and 117 CATH superfamilies. Rossman and Immunoglobulin folds were the among most represented in the dataset (Supplementary Figure1). These folds are amongst the most commonly occurring folds in the CATH database and a large proportion of UniProt sequences adopt one of these folds [62,63] (Supplementary Text10 and Supplementary Figure2 for details about model quality across superfamily). The remaining proteins and regions which were not assigned to CATH domains were scanned against Pfam FunFams. 416 domains belonging to 341 proteins were matched to 376 unique Pfam FunFams. A total of 496 regions of proteins (at least 50 residue long stretches) belonging to 332 proteins were designated as unassigned domains which did not map to CATH/Pfam domains. Some of these(41), could putatively be assigned to CATH superfamilies using more advanced sequence search strategies and structure comparisons using the predicted structure (see Supplementary Text12) [64,65].

### 2. Comparison of domain models generated using RoseTTAFold and AlphaFold

The quality of the models was assessed using the average predicted lDDT score [66] (Supplementary Text13 for details about lDDT). The range of lDDT scores for RoseTTAFold models was between 0 and 1, while those of AlphaFold were between 0 and 100. RoseTTAFold models with a score of 0.7 and AlphaFold models with a score of 70 were considered as good models, as recommended by the developers of these techniques. 304/309 (98%), 328/416 (79%) and 198/469 (42%) CATH, Pfam and unassigned domains respectively were good AlphaFold models (Figure 1). Whereas for the RoseTTAFold models 274/309 (89%), 240/416 (58%) and 79/469 (17%) CATH, Pfam and unassigned domains respectively were good models. 72% AlphaFold models and 58% RoseTTAFold models were of good quality.

**Figure 1-.**
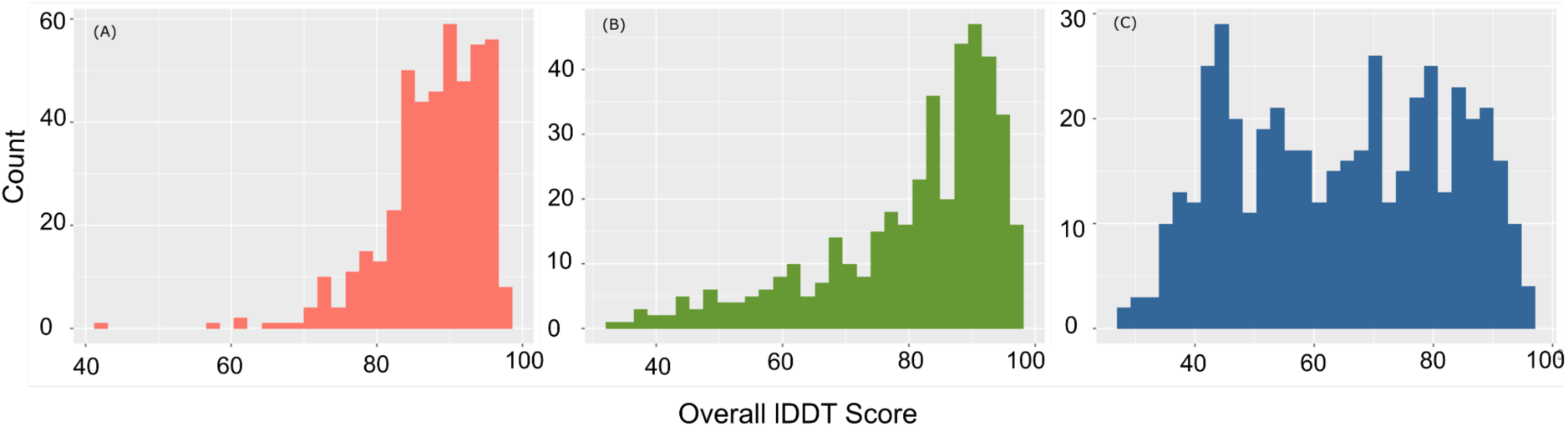
The model quality of AlphaFold models for A) CATH domains B) Pfam domains C) Unassigned domains.

We did not have access to the MSAs used to build the AlphaFold models, hence we analyzed the MSAs used to build the RoseTTAFold models. We calculated the Neff, DOPs, percent scorecons and number of taxons in the alignment to check the information content of the MSA [67] (Supplementary Text13 for more details). These measures quantify the diversity of the sequences in the alignment and not the quality of the alignment. Neff and the number of taxons in the alignment are higher for CATH models compared to the other two groups providing more coevolutionary information for those models (Supplementary Figure3). However, no clear correlation was found between Neff, DOPs, percent scorecons, number of taxons in the alignment and model quality (Supplementary Figure4). This trend is similar to that reported by AlphaFold and RosettaFold (Details in Supplementary Text2) [35,36].

Though the AlphaFold model quality was higher for 89% of the models compared to RoseTTAFold, 2, 27 and 45 CATH, Pfam and unassigned domains respectively had good RoseTTAFold models where the AlphaFold models were not of good quality (Figure 2). It should also be noted that the AlphaFold domains were extracted from the full proteins, however, RoseTTAFold models were built directly from the domain sequences. Residues in AlphaFold domains were modelled in the presence of the residues from other interacting domains, which was not the case for RoseTTAFold. Hence added constraints during modelling might have led to the better model quality of the AlphaFold models as compared to RoseTTAFold models.

**Figure 2-.**
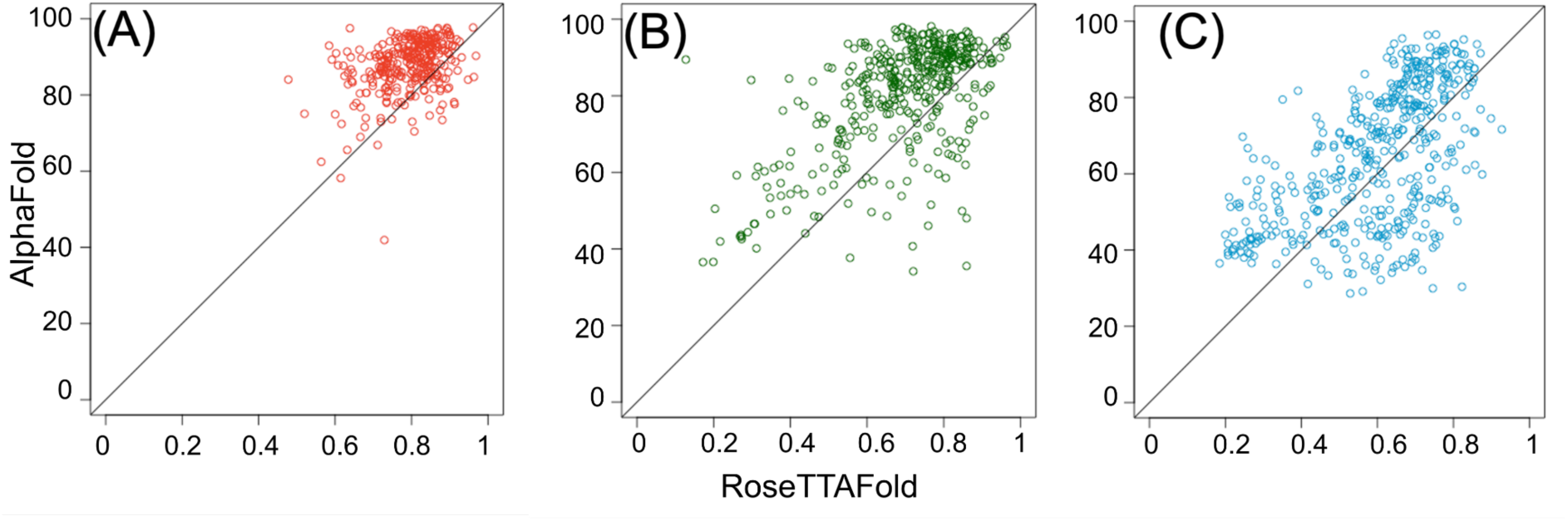
Model quality of AlphaFold models vs RoseTTAFold models for A) CATH domains B) Pfam domains C) Unassigned domains.

Around 88%, 74% and 64% of the CATH, Pfam and unassigned domains had an RMSD < 2Å between domains which had both good RoseTTAFold and AlphaFold models, indicating higher structural similarity for the CATH and Pfam domains for good quality models (Supplementary Figure5). In some cases, simple orientation changes of helices or sheets with respect to one another in RoseTTAFold domain models would help in better superimposition to those of their AlphaFold counterparts, hence reducing the RMSD. Some of the models, including regions of the C2 domain-containing protein 3 (Q4AC94) mapped onto CATH FunFams, showed RMSD < 1A between RoseTTAFold and AlphaFold models (Figure 3). As expected, the RMSD of the domains increased with a decrease in model quality (Figure 4). The correlation of RMSD with model quality is as expected and provides confidence in the techniques and their independence. The length of the model did not affect the model quality of AlphaFold models (Supplementary Figure6 and Supplementary Text11).

**Figure 3 -.**
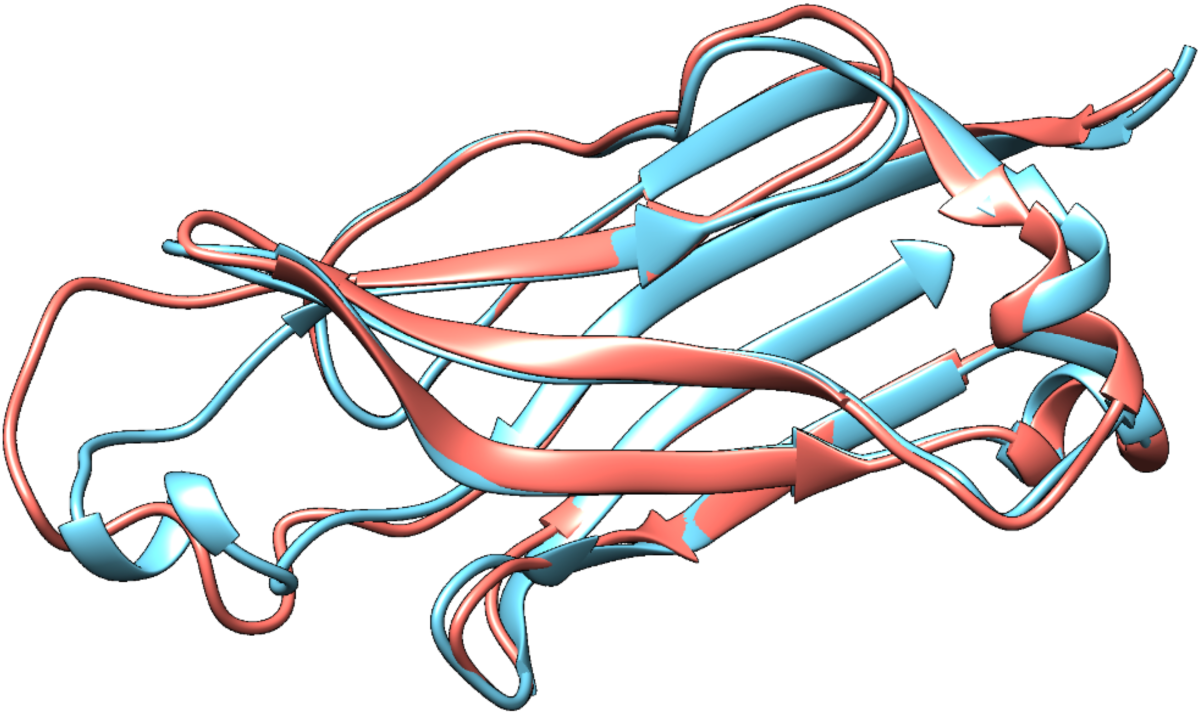
Ribbon model of superimposition of AlphaFold model (salmon) and RoseTTAFold model (sky blue) A) CATH domain of C2 domain-containing protein 3, having an RMSD of 1 Å. Both the models are of good quality.

**Figure 4 -.**
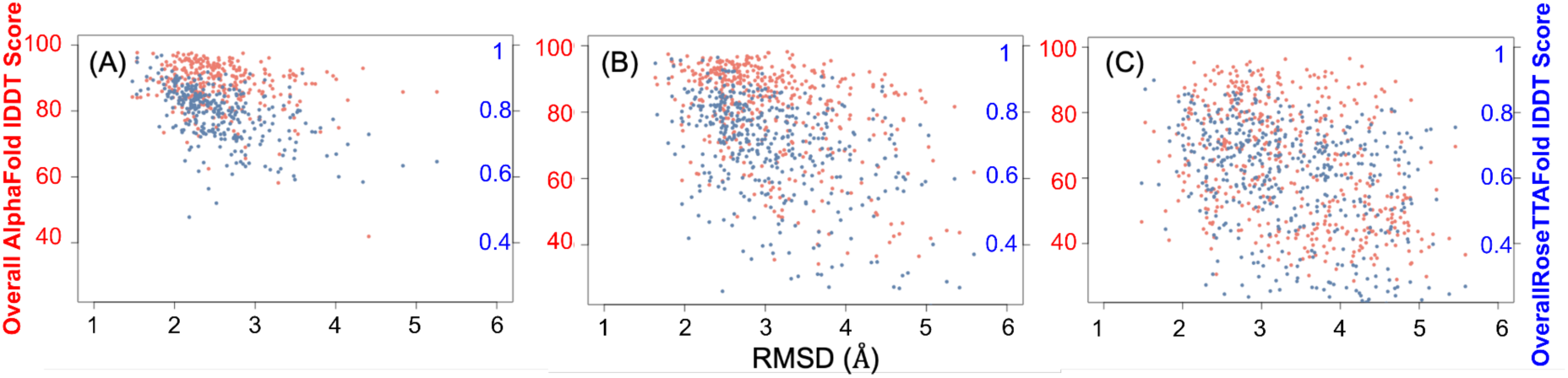
Distribution of model quality (AlphaFold in red, RoseTTAFold in blue) against the RMSD between the models for a) CATH domains b) Pfam domains and c) Unassigned domains

### 2. Disorder in protein domains

The probability of a residue in the sequence to be disordered for the CATH, Pfam and unassigned domains was calculated using IUPred2A [68]. We then calculated the percentage of the domain sequence that is disordered. For the CATH, Pfam and unassigned domains around 3, 15 and 38% of the sequences respectively had greater than 40% of its sequence disordered (Figure 5). This indicates that the CATH domains had much lower disorder compared to Pfam which in turn was much lower compared to the unassigned domains. Given the fact that the disordered regions lack proper structure, the model quality of those regions will be low, hence lowering the overall domain model quality. The percentage of the predicted disordered region in the sequences of the unassigned domains had an inverse correlation coefficient of 0.74 and 0.41 with model quality for the AlphaFold and RoseTTAFold models respectively (Supplementary Figure7), indicating that the model quality was lower because the sequences might be disordered. A low model quality for disordered regions provides higher confidence in the model quality scores.

**Figure 5 -.**
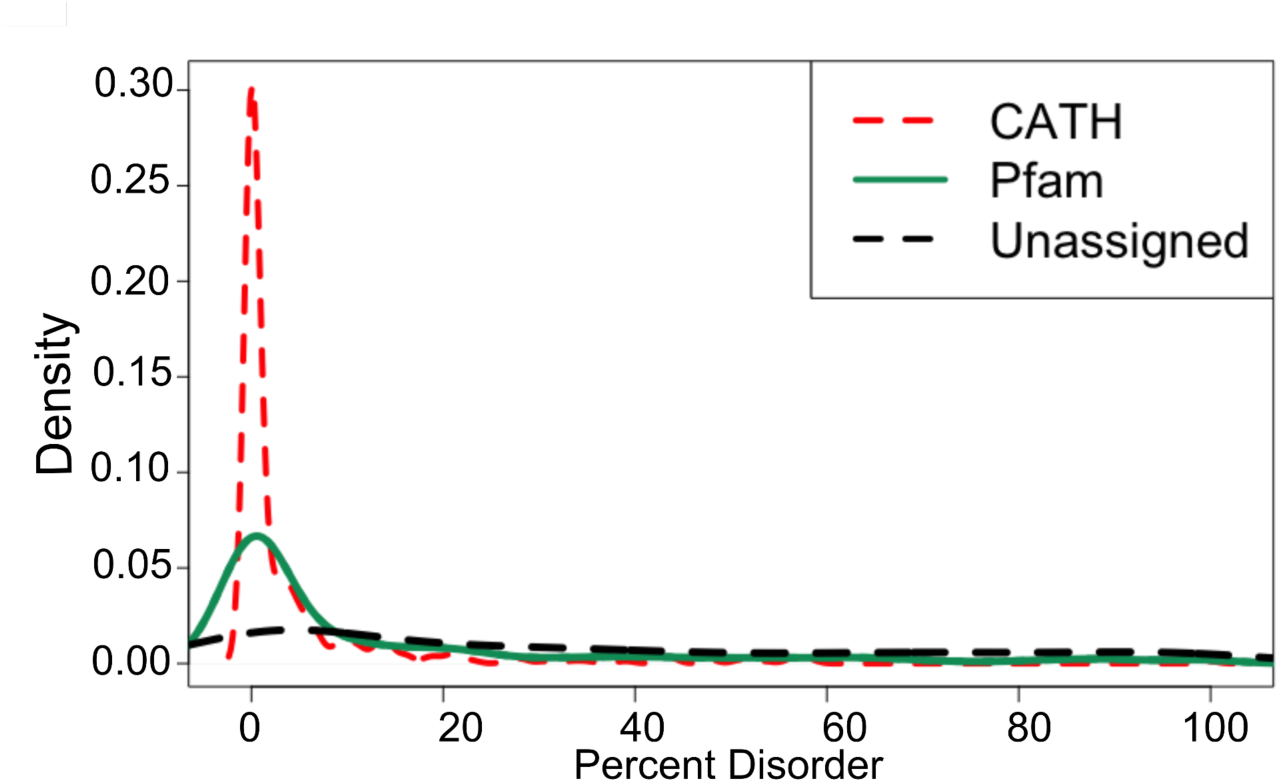
Density plot showing the distribution of percent disorder of the domain sequence for CATH (in red dashed line), Pfam (in green) and unassigned (in black dashed line) domains.

### 3. Characterization of disease-associated mutations

The total number of disease-associated missense mutations for our 553 modeled proteins in humsavar is 1730, corresponding to 1568 residue positions. Around 81% of the proteins in our dataset had 1 disease annotation for all the mutations. 40 of the proteins had more than 10 disease-associated mutations (Supplementary Figure8A) of which 19 proteins had a single disease annotation (Supplementary Figure8B) (Details in Supplementary Text3). About 78% of the residues involving mutations leading to the same disease are structural neighbours (within 5 Å), suggesting a similar mechanism. The disease annotations in our dataset were mostly nervous system disorders followed by musculoskeletal system disorders according to Human Disease Onology database [69] (Supplementary Figure9 and Supplementary Text3).

We checked if any of the mutations corresponded to gain of function (GOF)/loss of function (LOF) from GOF/LOF database [70] based on Human Gene Mutation database [71] and MAVEDB [72]. GOF/cancer mutations are usually clustered together in 3D space [73–75]. 13 out of 14 GOF mutations in the GOF/LOF database involve structural neighbours, while MAVEDB had no proteins from our dataset (Supplementary Text3).

We examined the relative abundance of the amino acids that the residues mutated into against proteins in the SwissProt51 database, using Composition Profiler [76]. We noticed an enrichment of Cys, His, Pro, Arg, Ser and Trp with p-value <0.05 (Supplementary Figure10 and Supplementary Table1). These amino acids are either neutral polar or basic polar. In addition, Trp is a bulky amino acid and mutation into Trp can structurally de-stabilize the protein. Mutation into Cys might introduce unfavourable disulphide bonds and Pro can disrupt helical structures. These propensities are similar to a much larger amino acid mutation database study [77] for human disease mutations.

### 4. Explaining disease-associated mutations

We checked if these mutations were on or near a predicted ligand binding site (by P2Rank) or protein-protein interface (by meta-PPISP) or are conserved (by scorecons) in the FunFam or superfamily. Though there are a lot of servers and tools for predicting ligand binding sites and protein-protein interface which we recently reviewed [41]. We used meta-PPISP and P2Rank based on availability, ease of installation, ability to run the scripts on all the protein domains and usage in recent studies (Details about choice of tools in Supplementary Text4) [24,37,78– 82].

We considered the residues within a 5 Å radius (heavy atom distance) (see supplementary Text4 for explanation of this threshold) [83–85] of a predicted functional site because mutations in those can lead to change in the environment of the functional site residue, potentially leading to modification or loss of activity. We limited our analysis to the structure of the wild type protein as we were unsure of the ability of the techniques to build the mutant models. We also calculated the effect of the mutations on the stability of the protein using FoldX and DynaMut2 and pathogenicity of mutations using MutPred2.

Out of the 1568 disease associated residue positions, 1317 residue positions (corresponding to 1389 mutations) are in high confidence AlphaFold modelled domains and residues (lDDT>70). We do not analyse residues with lDDT<70 as those are low confidence regions of the models and hence should be treated with caution. For these domains, 122 CATH (585 residue positions), 143 Pfam (508 residue positions) and 61 unassigned domains (224 residue positions) had disease-associated mutations.

796 of these disease-associated residue positions (corresponding to 919 mutations) are either in/near a predicted functional site (Figure 6). In addition, 272 mutations (corresponding to 227 residue positions) were predicted to be destabilising using FoldX (Figure 5). Other than these, 8 mutations (corresponding to 8 residue positions) were predicted to be pathogenic. Of all the disease-associated residue positions, 488 were near the predicted protein-protein interface, 163 near predicted ligand binding sites, 473 near conserved sites. 816 mutations (corresponding to 689 residue positions) were predicted to be destabilising by FoldX (ddG>1) (Supplementary Figure11) (see Supplementary Text4 for explanation of this threshold). We used two independent methods, FoldX and DynaMut2 to predict the free energy change of mutation. We found that for about 95% of the mutations predicted to have an impact by FoldX, DynaMut2 [86] also predicted impact, giving higher confidence in the predictions (Supplementary Figure11). We have not used the computed ddG value to quantify the impact of the mutation as such computed values might not correspond to actual ddG values. 899 mutations (corresponding to 844 residue positions) were predicted as pathogenic using MutPred2 (Figure 9A). Of these, 638 mutations (corresponding to 629 proteins) were in the functional sites. In total, this suggests that we can provide some rationale for 78% of the disease-associated mutations in the well modelled regions.

**Figure 6 -.**
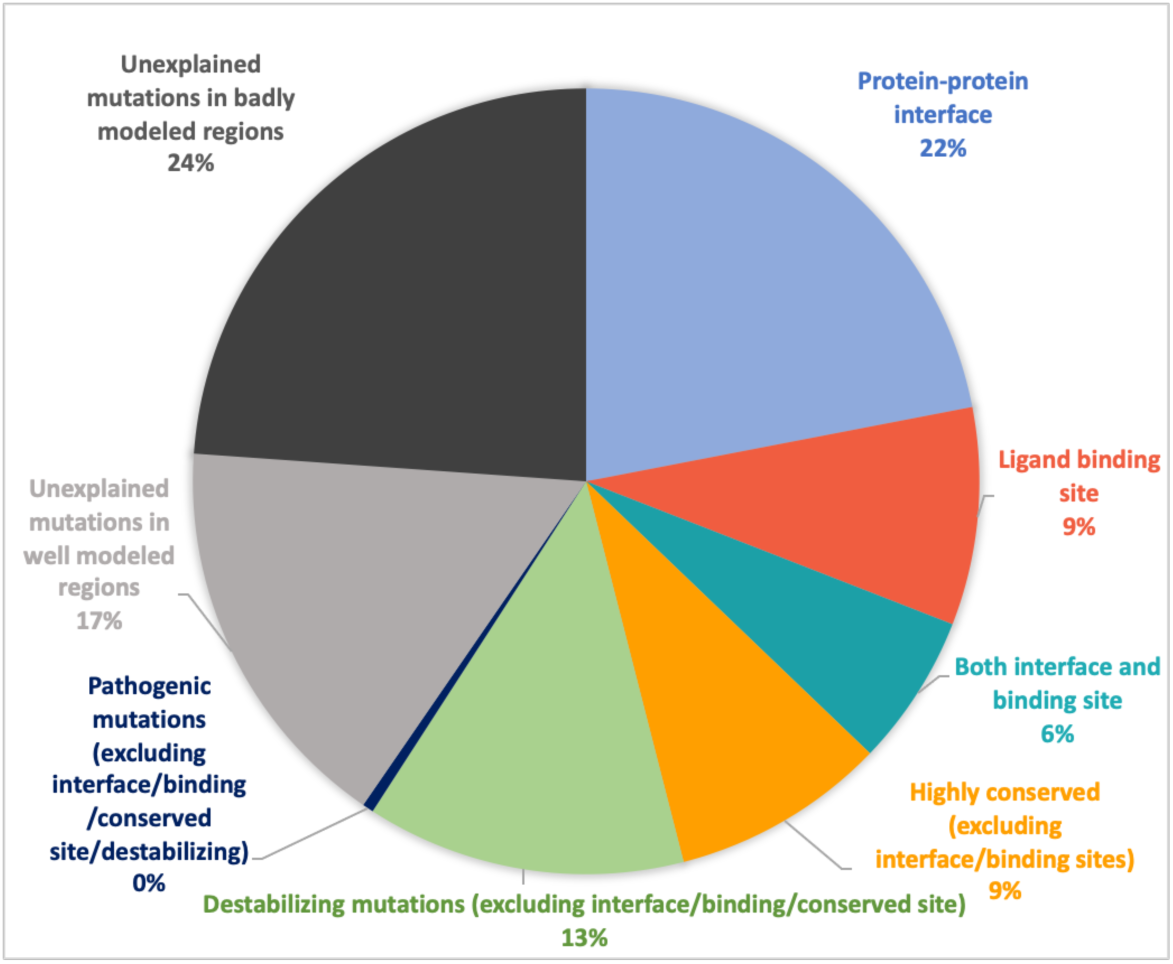
Percentage of disease-associated mutations that are predicted to be in/near a ligand binding/protein-protein interface/conserved site in the AlphaFold domain models. Some of these mutations are predicted to be destabilizing or pathogenic.

We calculated the solvent accessibility of residues whose mutations lead to disease and compared them against the distribution of the solvent accessibility of residues that make up the entire domain (Supplementary Figure19). We noticed that 42% (547 residues) of the residues that lead to disease were buried (Residues with solvent accessibility < 20 were considered buried [87]) (Figure 9). This is much higher than the 21% of residues buried in the entire domain. A large number of the disease associated mutations are buried, which could cause destabilisation of the protein. Out of the 547 buried residues, 428 residues had mutations that were predicted destabilising according to FoldX.

When we consider whether mutations fall on overlapping functional sites, we observe that 108 mutations were predicted as both near a ligand binding site and a protein-protein interface, 101 mutations either near a conserved residue or ligand binding site, 221 mutations were either near an interface or conserved site (Figure 7). Around 44% of these mutations have multiple evidence for it lying on/near a functional site, which provides higher confidence in these sites having functional significance.

**Figure 7 -.**
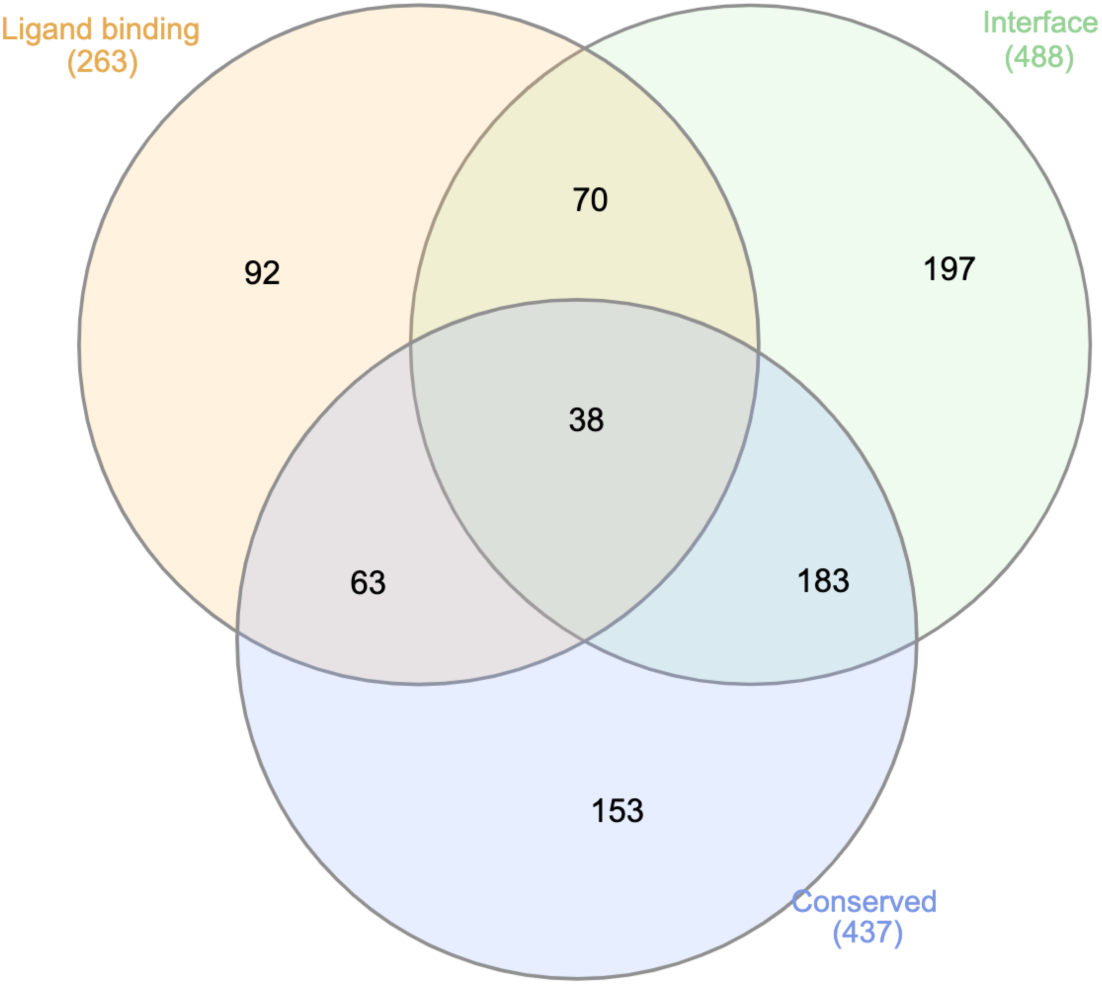
The overlap between the number of disease-associated mutation sites that are either predicted to be on/near a ligand binding/protein-protein interface/conserved for the AlphaFold domain models. Figure generated using InteractiVenn [120].

In addition, RoseTTAFold helped to find 93 additional mutations (in 44 proteins) near functional domain regions; which could not be explained using AlphaFold. The AlphaFold and RosettaFold models were very structurally similar to each other with an RMSD of 1.7+/- 0.6 Å (Supplementary Figure12). Small changes in orientations of the parts of the models led to low quality for AlphaFold model or prediction of functional site or larger distance of the functional site from mutations (Supplementary Figure13 and Supplementary Text9). There was an overlap of 492 mutations predicted to be a functional site between the predictions derived from RoseTTAFold and AlphaFold models, providing higher confidence in these predictions.

Mutations in residue numbers 32,33,233,236 and 412 in transmembrane helices of putative sodium-coupled neutral amino acid transporter 8 (Uniprot A6NNN8; Gene SLC38A8) lead to foveal hypoplasia 2. Other than residue 412, the others are spatially close to each other and are predicted either to be in/near a ligand binding site. In addition, residue 233 is also predicted to be near a conserved site (Figure 8A). Mutations in Vitamin K Epoxide Reductase Complex submit 1 (Uniprot Q9BQB6; Gene VKORC1) leads to Coumarin resistance, which lie on/near the predicted ligand binding sites and interface residues (Figure 8B). In another example, the immediate early response-3 interacting protein-1 (Uniprot Q9Y5U9, Gene IER3IP1) RoseTTAFold models had acceptable model quality whilst the AlphaFold model was of low quality or the residue of interest was of low quality. Mutations in residue numbers, 21 and 78 (associated with microcephaly, epilepsy, and diabetes syndrome (MEDS)) are on the predicted protein-protein interface site (Figure 8C). In certain cases such as mutations in Beta-1,4 N-acetylgalactosaminyltransferase 1 (Q00973) and Feline leukemia virus subgroup C receptor-related protein 1 (Q9Y5Y0) could only be explained by large ddG (>2.5) (Supplementary Figure14 and Supplementary Text8).

**Figure 8 -.**
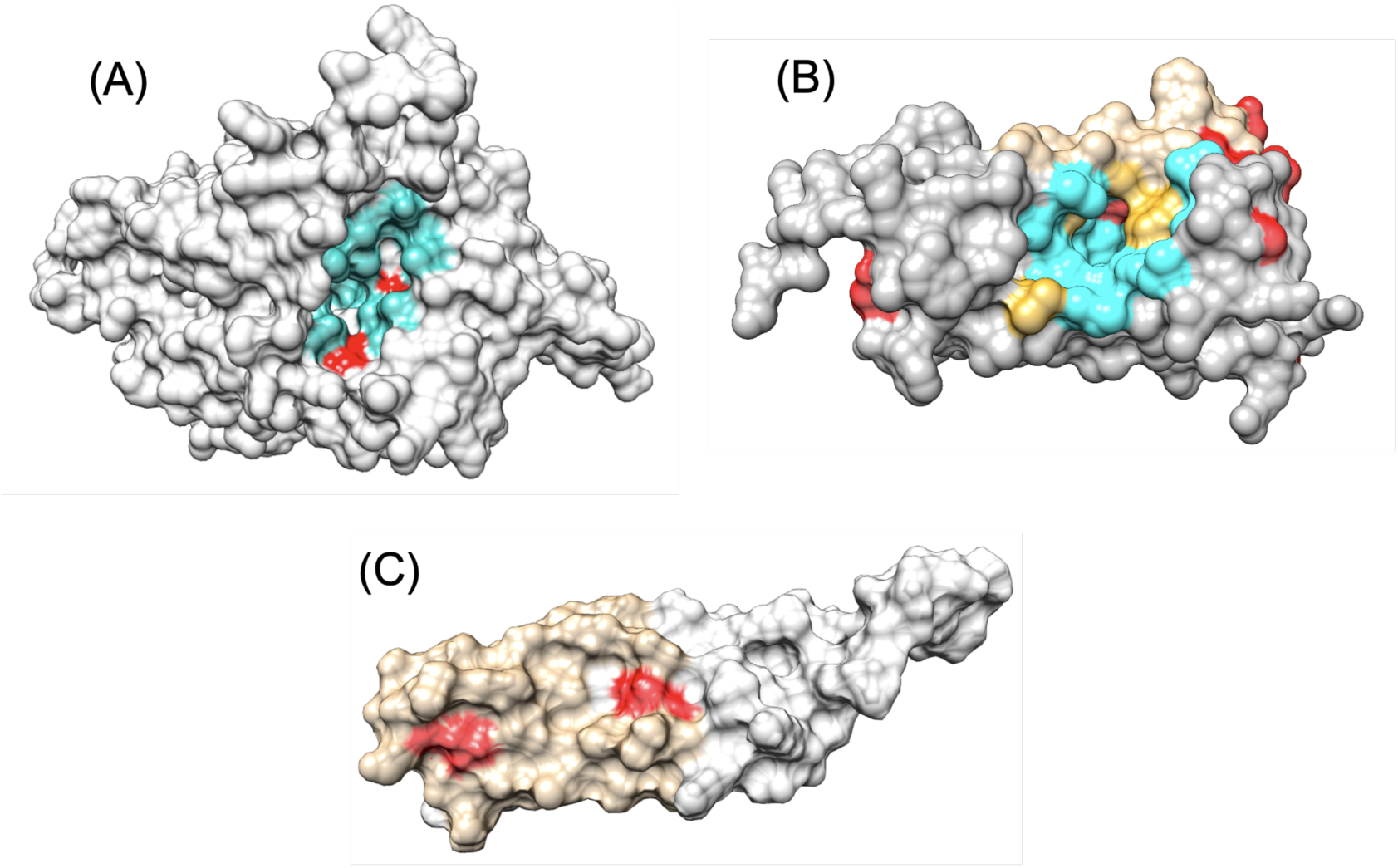
A) AlphaFold models in grey surface representations for Putative sodium-coupled neutral amino acid transporter 8, the residues in cyan are the predicted ligand binding residues and in red are the disease-associated mutations. B) AlphaFold models in grey surface representations for Vitamin K Epoxide Reductase Complex submit 1, the residues in cyan are the predicted ligand binding residues, tan are the predicted interface residues, golden are the residues predicted both as ligand binding and interface, and in red are the disease-associated mutations. C) RoseTTAFold model of early response-3 interacting protein-1 in grey surface representation. The predicted interface is in tan and the disease-associated mutations are in red.

We also looked at the PANTHER protein classes, to check if different functional sites were enriched in the different classes (Supplementary Figure17). Given our limited dataset, we could not compare the various properties across all the protein classes (see analysis of 3 PATHER classes in Supplementary Text5), however a detailed study across all the classes has previously been done for a larger dataset of human proteins, by Iqbal and coworkers [88].

### 5. Comparison of disease associated mutations and polymorphisms

In order to compare the disease associated mutations with that of likely neutral variants (polymorphisms), we checked the ddG of mutation, proximity to predicted functional sites, pathogenicity and solvent accessibility of these polymorphisms in the AlphaFold models. The number of residues with polymorphisms were 998 (see Supplementary Text6). We see that a larger percentage of disease associated mutations were closer to functional site, buried, destabilising and pathogenic as compared to polymorphisms (Table 1 and Figure 9). The distributions of the pathogenicity scores, relative solvent accessibility, ddG for the polymorphisms and disease associated mutations are significantly different (p-value < 2.2e-16) when compared using the Mann-Whitney Wilcox test [89]. When we compared the relative abundance of amino acids in deleterious mutations (compared to amino acids in polymorphisms) we noticed that Cys, Pro, Arg, Tyr, Trp are more predominant in deleterious mutations compared to polymorphisms (Supplementary Figure16) (Results Section 3 for effect of these mutations on protein function/structure).

**Figure 9 -.**
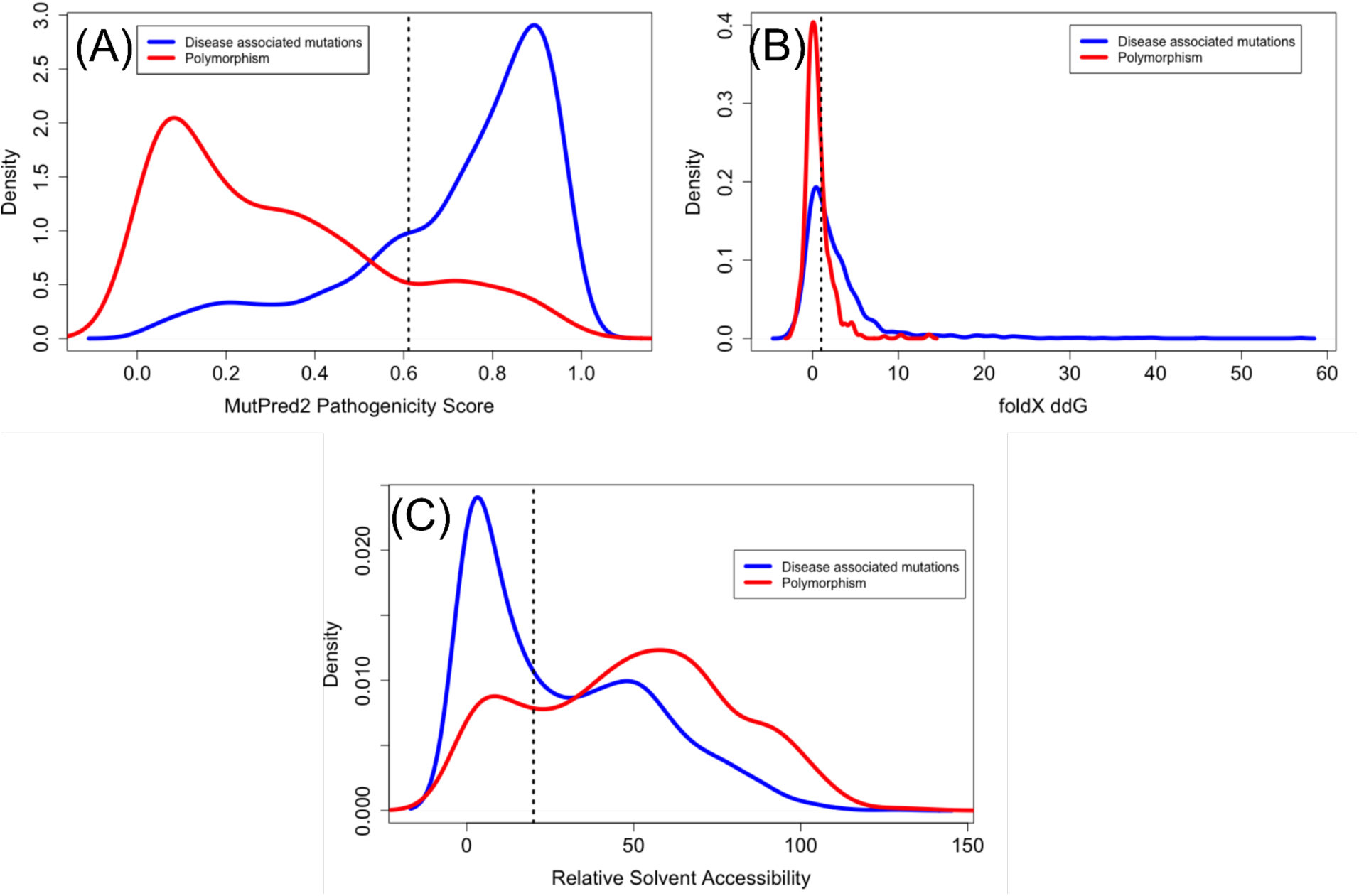
Density plots showing the distribution of A) MutPred2 predicted pathogenicity scores B) FoldX predicted ddG values C) relative solvent accessibility of residues that have disease associated mutations in blue and polymorphism in red. The dotted horizontal lines show the cutoff beyond which the mutations are predicted as pathogenic (>0.611), destabilizing (>1) and buried (<20)

**Figure 10 -.**
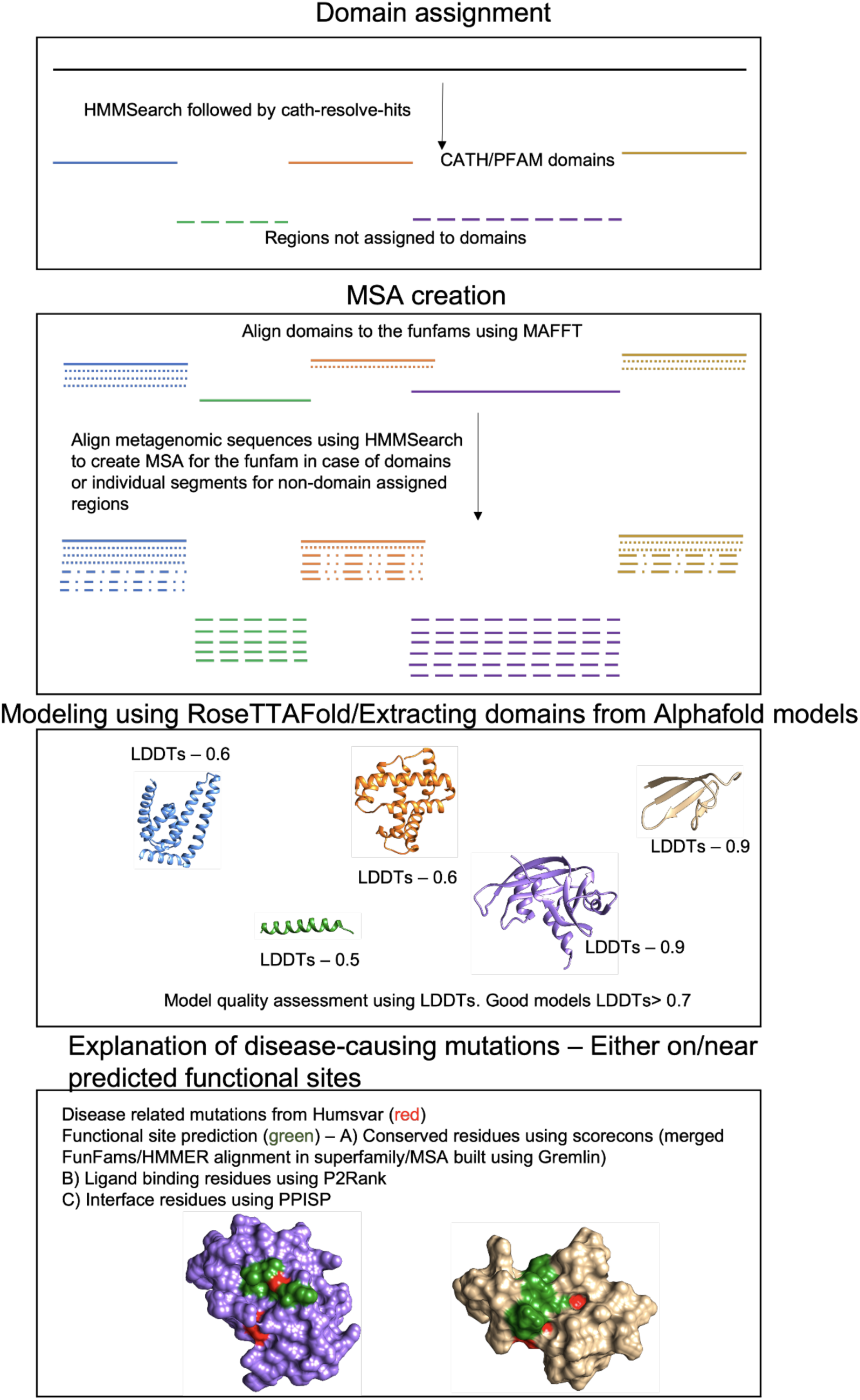
Schematic explaining the methodology to model protein domains and explain disease-associated mutations.

**Table 1-.**
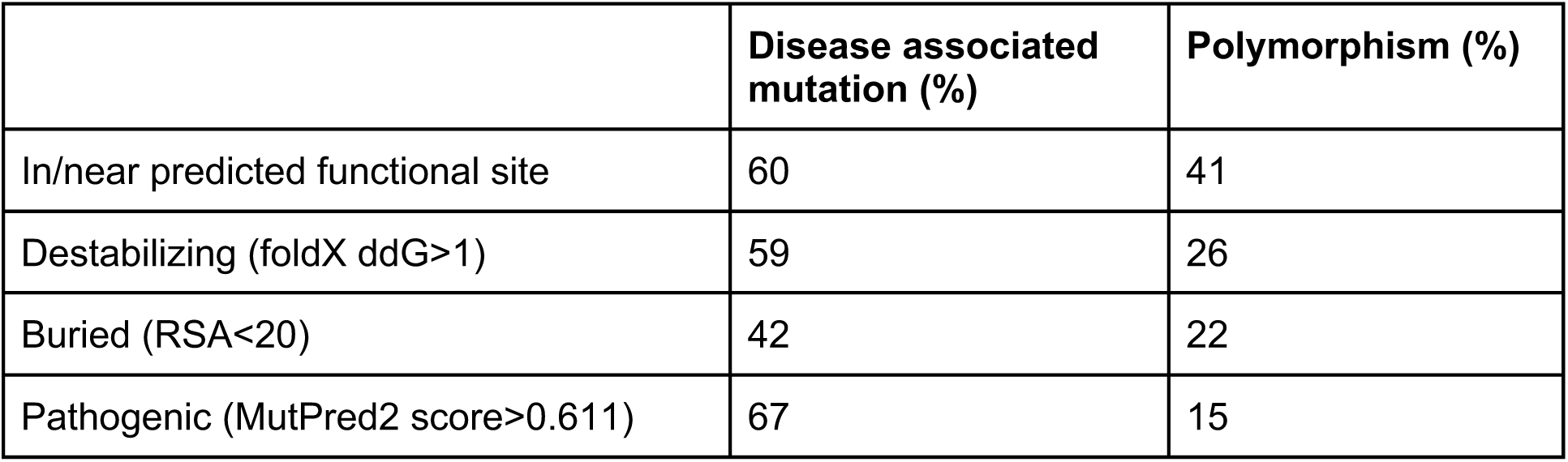
Table showing the percentage of residues that are near functional site, buried, destabilizing and pathogenic for a residue that has a disease associated mutation compared to those residues with polymorphism.

|  | **Disease associated mutation (%)** | **Polymorphism (%)** |
| --- | --- | --- |
| In/near predicted functional site | 60 | 41 |
| Destabilizing (foldX ddG>1) | 59 | 26 |
| Buried (RSA<20) | 42 | 22 |
| Pathogenic (MutPred2 score>0.611) | 67 | 15 |

## Discussion

With the current improved state of art computational structural predictors like RoseTTAFold and AlphaFold, we can model regions of proteins without any known homologues for about 20% of the residues in humans [90]. In this study, we modelled the disease-associated human proteins without known homologues (calculated using BLAST) using RoseTTAFold and AlphaFold. We used these models to check if the disease-associated mutations were near predicted functional residues (i.e. ligand binding site/protein-protein interface/conserved residues). Some of these disease-associated mutations were also predicted to be structurally destabilizing or pathogenic.

For our analyses we used a method BLAST; most commonly used by biologists to detect structural homologues in the PDB. However, in principle more sophisticated sequence search tools could be used and biologists may seek to employ MODELLER [26] or similar tools. We checked using MODELLER and this resulted in ∼6% of the domains (at least 50% of the protein sequence with at least 30% sequence identity) having homologues in the PDB, giving no substantial changes to the trends (Supplementary Figure18 and Supplementary Table3).

The AlphaFold models (70%) were largely of better quality when compared to those of RoseTTAFold models (50%). However, there were a total of 74 models where RoseTTAFold produced a good quality model whereas AlphaFold had a low quality model. We should keep in mind that model quality calculations for the two techniques were independent of each other and no consensus model quality assessment tool was used. To date, there has been no evaluation of the model quality estimates of AlphaFold and RoseTTAFold. However, optimising model quality cut-offs for such consensus estimators will involve benchmarking and is out of the scope of the current study. The high confidence models (where both the techniques provide consensus structures) could be used to predict/design inhibitors/drugs against them [91], predict off-target effects of putative drugs [92], explain the functioning of proteins, the impact of variations [93,94] or model protein complexes [95].

The CATH domains were modelled better compared to domains that could only be assigned Pfam or not assigned to either because of lower disorder and greater diversity in the alignments used for modelling. With the ease of sequencing and large scale sequencing projects, we expect to further increase the repertoire and quality of protein sequences in the near future. With the increased number of sequences, the model quality of the Pfam and unassigned domains will hopefully improve.

Functional sites such as ligand binding sites, interface residues, allosteric sites, post-translational modification sites etc are more conserved compared to the rest of the protein, hence mutation of these conserved sites and their neighbours might lead to a loss/modification in protein function. We considered residues close to the predicted functional site because these residues can have an effect on the structure and functioning of the functional site residues and hence changes to them might also alter the functioning of the protein. Also, previous studies have shown that in a large number of cases, the disease-associated mutations are near functional sites [77,96]. We assume that the neighbourhood of the functional sites will be well predicted at this level of model quality even if the functional residues are not always correct. Interface residues and ligand binding sites have been shown to be highly conserved [97]. In our dataset, 38% of the disease-associated ligand binding sites and 45% of the protein-protein interfaces are conserved, hence providing higher confidence that these mutations could impact structure or function. Also, around 59% of these mutations were predicted to be structurally destabilising (based on FoldX), which might also impact structure and function. 85% of these disease associated mutations in the buried residues were predicted to be destabilising. Around 67% of the disease associated mutations were also predicted to be pathogenic. We also had 37% of the mutations being predicted near functional sites using both AlphaFold and RoseTTAFold models, which provide additional confidence in those predictions. We were able to explain 78% of the disease-associated mutations in the good quality regions. Given the fact that the models built using the state of art tools AlphaFold and RoseTTAFold might still have inaccuracies, cases, where the structure predicted using multiple techniques, are similar provide higher confidence in the prediction accuracies. We also compared the disease associated mutations with the polymorphisms and found a larger proportion of disease associated mutations were more buried, destabilising, pathogenic and had a larger percentage closer to predicted functional sites.

The fact that a large number of disease causing mutations could be explained using the AlphaFold models, further validates the model quality as we would expect poor models to introduce noise to our analyses. Predicting whether mutations are likely to be disease associated remains one of the most challenging tasks in bioinformatics, despite the improvements brought about by deep learning. This is because multiple factors can lead to a disease phenotype such as stability of the protein, change in binding affinity, expression levels of the protein, minimum concentration of the proteins required for its functioning etc. Our study can help in providing insights on the structural and functional features that are frequently disrupted by disease-causing mutations.

In summary, the strategy of using good quality protein structures to predict functional sites can help explain the mechanism of disease-associated mutations. Similar strategies can be used on good quality models of the entire human proteome in the future to explain disease associated mutations.

## Methods

### 1. Dataset of disease associated human proteins to model

#### a. Curation of disease associated human proteins

The VarSite database [53] contains 4444 human proteins which are disease-associated as annotated in Uniprot or ClinVar. Out of these, 553 human disease proteins are without any structural homologues (match determined by running the sequence with BLAST [98] against all PDB [52] sequences with evalue cutoff of 1e-06). We used PANTHER to study the GO enrichment of these proteins. In addition, we divided the proteins into the 24 PANTHER classes (Supplementary Table2) to study the properties of each group.

#### b. Assignment of domain boundaries to disease-associated human proteins without structural homologues

All the 553 proteins were scanned against CATH-FunFams v4.3 (212,872 HMMs) using HMMER3 [99] with an e-value cut off of 1e-03 to assign domains. The domain boundaries were resolved using cath-resolve-hits [100] with a bitscore cut-off of 25 and coverage of 80%. The proteins/regions of proteins that were not assigned to CATH domain boundaries were scanned against the Pfam32-FunFams (102,712 HMMs) using the same protocol. The regions not belonging to domains identified by scans against CATH/Pfam FunFams, were modelled separately. These regions have been referred to as unassigned domains.

### 2. Modeling the proteins using RoseTTAFold

#### a) Multiple sequence alignment (MSA) generation for protein domains

The domains that were assigned to the CATH/Pfam FunFams were aligned to the FunFams using MAFFT (version 7.471) [101] using the default options. The resulting seed alignments were enriched with additional homologs by performing iterative sequence *hhblits* [102] searches against uniclust30 (version UniRef30_2020_01) [103] and BFD [104,105] databases as outlined by Anischenko and colleagues [106] (Details in Supplementary Text14).

#### b) Assessment of diversity of MSA

The diversity of the alignment was accessed using 4 parameters - Neff, DOPs (Diversity of Positions), percent scorecons and number of taxon IDs in the alignment (Details in Supplementary Text13). DOPs and scorecons were calculated using <u>Cathy</u>.

#### c) Modeling of the protein domains

Using multiple sequence alignments from Section 2a above, structure templates were identified by *hhsearch* [102] searches against the PDB100 database and 10 best scoring templates were selected. The identified templates along with the multiple sequence alignment were then used as inputs to RoseTTAFold [36] to predict residue-residue distances and orientations followed by the PyRosetta-based [107] 3D structure reconstruction protocol from trRosetta [33]. 15 structure models were generated and subsequently rescored according to lDDT scores [66] predicted by DeepAccNet [108]. As revealed by the results of the CASP14 experiment [106], rescoring the pool of the PyRosetta-generated models with DeepAccNet brings in an additional 1-2 GDT_TS score [109] units improvement in the model accuracy, beyond what can be achieved from picking models by Rosetta energy [110]. The model with the highest predicted global lDDT score was selected for further analysis.

### 3. Creating domains from AlphaFold models

The domains were extracted from the AlphaFold models downloaded from the AlphaFold DataBase portal. The domain boundaries were the same as obtained by the CATH/Pfam scans or unassigned domains as mentioned in Section 1. The AlphaFold domains were superimposed on the domains built using RoseTTAFold using GESAMT [111].

### 4. Scoring the local and global quality of the models

The model quality for both RoseTTAFold and AlphaFold models were calculated using the predicted lDDT scores; which is commonly used by CAPRI [66,112] (Details in suppplementary S9). The global quality of the model is calculated as the average of the local lDDT scores of the constituent residues.

### 5. Disorder prediction on the domain sequences

The disorder percentage was calculated for the unassigned regions using IUPred2A [68] using default parameters. A residue was considered disordered if the prediction probability was greater than 0.5.

### 6. Prediction of functional sites on protein domains and known mutational sites

The human mutations were identified from humsavar [23] as downloaded on 2nd December 2020. The mutations were classified as disease (used for disease associated mutations), polymorphism and unclassified (Supplementary Text7 for current nomenclature). The conserved positions were identified as positions in the MSA with a high scorecons value (>=0.8) or position of > 0.65 scorecon value near (<5 Å) a position of high scorecon value (>0.8) (Details about MSA generation for identification of conserved residues in Supplementary Text14). The neighbourhood of the predicted functional residues were identified using the implementation of the cell list algorithm [113–116] (Details about choice of algorithm in Supplementary Text4).

The ligand binding sites were predicted using P2Rank [117], with default parameters. Positions were annotated as ligand binding if the probability of the ligand binding was >=0.5 (intuitively chosen). The protein-protein interaction sites were predicted using the meta-PPISP webserver [118]. The mutation was predicted to affect the functioning of the protein if the site was either on or near (within 5 Å) of a predicted functional site by P2Rank/meta-PPISP/highly conserved position. We only considered predictions for those sites where the entire domain had a good model quality score and the local quality score of the mutated residue was good (lDDT score of 70 for AlphaFold models and 0.7 for RoseTTAFold models).

We calculated the relative solvent accessibility of the residues using NACCESS [119]. For each of these sites, the effect of mutation on the stability of the protein was identified using FoldX [43]. In order to be stringent; ddG>1 was chosen as destabilising. In addition to FoldX, the ddG of the mutation was also calculated using DynaMut2 API [86]. This technique predicts a mutation as destabilising if ddG<0. The likely pathogenic impact of the mutations was predicted using the stand-alone version of MutPred2 [47] using default parameters (Details about choice of tool in Supplementary Text4).

## Data availability

We have also tabulated all the disease-associated positions, their predicted functional site impacts, ddG of mutations, pathogenicity scores and other characteristics <u>here</u>. The table also lists the known or predicted functions of the proteins given by the experimental GO terms for the FunFam in which the proteins are classified. All the data and the models can be accessed <u>here</u>.

## Conflict of Interest

None.

## Acknowledgements

The following people acknowledge the Biotechnology and Biological Sciences Research Council for their funding: NS [BB/S020144/1], NB [BB/R009597/1], IS [BB/R014892/1] and SV[BB/S017135/1]. IA is supported by National Science Foundation Award # DBI 1937533 and DB is supported by Audacious Project at the Institute for Protein Design. We would like to acknowledge Roman Laskowski for help with VarSite data and Paul Ashfold for help with disease datasets.

## Author Contributions

**Conceptualization – CO, DB, SV**

**Methodology - CO, DB, SV, NS, IA, IS**

**Software – NS, IA, NB, IS**

**Validation – NS, IA**

**Formal Analysis – NS, IA, NB**

**Investigation – NS, IA, NB**

**Resources – CO, DB**

**Data Curation – NS, IA**

**Writing – Original Draft - NS**

**Writing – Review & Editing – NS, IA, NB, SV, CO**

**Visualization - NS**

**Supervision - CO, DB, SV, IS**

**Project Administration - CO, DB, SV**

**Funding Acquisition - CO, DB, SV**

## Notes

https://zenodo.org/record/6359971#.YkLfyC8w2_w

### Competing Interest Statement

The authors have declared no competing interest.

## References

1. Coleman CS, Stanley BA, Pegg AE. Effect of mutations at active site residues on the activity of ornithine decarboxylase and its inhibition by active site-directed irreversible inhibitors. J. Biol. Chem. 1993; 268:24572–24579

2. Joshi S, Virdi S, Etard C, et al. Mutation of a serine near the catalytic site of the choline acetyltransferase a gene almost completely abolishes motility of the zebrafish embryo. PLOS ONE 2018; 13:e0207747

3. Ricatti J, Acquasaliente L, Ribaudo G, et al. Effects of point mutations in the binding pocket of the mouse major urinary protein MUP20 on ligand affinity and specificity. Sci. Rep. 2019; 9:300

4. Lee B, Thirunavukkarasu K, Zhou L, et al. Missense mutations abolishing DNA binding of the osteoblast-specific transcription factor OSF2/CBFA1 in cleidocranial dysplasia. Nat. Genet. 1997; 16:307–310

5. Jubb HC, Pandurangan AP, Turner MA, et al. Mutations at protein-protein interfaces: Small changes over big surfaces have large impacts on human health. Prog. Biophys. Mol. Biol. 2017; 128:3–13

6. Cheng F, Zhao J, Wang Y, et al. Comprehensive characterization of protein–protein interactions perturbed by disease mutations. Nat. Genet. 2021; 53:342–353

7. Tyukhtenko S, Rajarshi G, Karageorgos I, et al. Effects of Distal Mutations on the Structure, Dynamics and Catalysis of Human Monoacylglycerol Lipase. Sci. Rep. 2018; 8:1719

8. Anfinsen CB. Principles that Govern the Folding of Protein Chains. Science 1973; 181:223–230

9. Soto C. Unfolding the role of protein misfolding in neurodegenerative diseases. Nat. Rev. Neurosci. 2003; 4:49–60

10. Baiardi S, Rossi M, Capellari S, et al. Recent advances in the histo-molecular pathology of human prion disease: Histo-molecular pathology of human prion disease. Brain Pathol. 2019; 29:278–300

11. Aznaourova M, Schmerer N, Schmeck B, et al. Disease-Causing Mutations and Rearrangements in Long Non-coding RNA Gene Loci. Front. Genet. 2020; 11:527484

12. Tan H. Somatic mutation in noncoding regions: The sound of silence. EBioMedicine 2020; 61:103084

13. Scacheri CA, Scacheri PC. Mutations in the noncoding genome. Curr. Opin. Pediatr. 2015; 27:659–664

14. Elliott K, Larsson E. Non-coding driver mutations in human cancer. Nat. Rev. Cancer 2021; 21:500–509

15. Smigielski EM, Sirotkin K, Ward M, et al. dbSNP: a database of single nucleotide polymorphisms. Nucleic Acids Res. 2000; 28:352–355

16. Fairley S, Lowy-Gallego E, Perry E, et al. The International Genome Sample Resource (IGSR) collection of open human genomic variation resources. Nucleic Acids Res. 2020; 48:D941–D947

17. Landrum MJ, Chitipiralla S, Brown GR, et al. ClinVar: improvements to accessing data. Nucleic Acids Res. 2020; 48:D835–D844

18. Forbes SA, Beare D, Boutselakis H, et al. COSMIC: somatic cancer genetics at high-resolution. Nucleic Acids Res. 2017; 45:D777–D783

19. Wang T, Ruan S, Zhao X, et al. OncoVar: an integrated database and analysis platform for oncogenic driver variants in cancers. Nucleic Acids Res. 2021; 49:D1289–D1301

20. Ainscough BJ, Griffith M, Coffman AC, et al. DoCM: a database of curated mutations in cancer. Nat. Methods 2016; 13:806–807

21. Stenberg KAE, Riikonen PT, Vihinen M. KinMutBase, a database of human disease-causing protein kinase mutations. Nucleic Acids Res. 1999; 27:362–364

22. Krassowski M, Paczkowska M, Cullion K, et al. ActiveDriverDB: human disease mutations and genome variation in post-translational modification sites of proteins. Nucleic Acids Res. 2018; 46:D901–D910

23. The UniProt Consortium, Bateman A, Martin M-J, et al. UniProt: the universal protein knowledgebase in 2021. Nucleic Acids Res. 2021; 49:D480–D489

24. PDBe-KB consortium, Varadi M, Berrisford J, et al. PDBe-KB: a community-driven resource for structural and functional annotations. Nucleic Acids Res. 2020; 48:D344–D353

25. Pei J, Grishin NV. The DBSAV Database: Predicting Deleteriousness of Single Amino Acid Variations in the Human Proteome. J. Mol. Biol. 2021; 433:166915

26. Šali A, Blundell TL. Comparative Protein Modelling by Satisfaction of Spatial Restraints. J. Mol. Biol. 1993; 234:779–815

27. Webb B, Sali A. Comparative Protein Structure Modeling Using MODELLER. Curr. Protoc. Bioinforma. 2016; 54:

28. Waterhouse A, Bertoni M, Bienert S, et al. SWISS-MODEL: homology modelling of protein structures and complexes. Nucleic Acids Res. 2018; 46:W296–W303

29. Rohl CA, Strauss CEM, Misura KMS, et al. Protein Structure Prediction Using Rosetta. Methods Enzymol. 2004; 383:66–93

30. Roy A, Kucukural A, Zhang Y. I-TASSER: a unified platform for automated protein structure and function prediction. Nat. Protoc. 2010; 5:725–738

31. Xu J. Distance-based protein folding powered by deep learning. Proc. Natl. Acad. Sci. 2019; 116:16856–16865

32. Greener JG, Kandathil SM, Jones DT. Deep learning extends de novo protein modelling coverage of genomes using iteratively predicted structural constraints. Nat. Commun. 2019; 10:3977

33. Anishchenko I, Ovchinnikov S, Kamisetty H, et al. Origins of coevolution between residues distant in protein 3D structures. Proc. Natl. Acad. Sci. 2017; 114:9122–9127

34. Senior AW, Evans R, Jumper J, et al. Improved protein structure prediction using potentials from deep learning. Nature 2020; 577:706–710

35. Jumper J, Evans R, Pritzel A, et al. Highly accurate protein structure prediction with AlphaFold. Nature 2021; 596:583–589

36. Baek M, DiMaio F, Anishchenko I, et al. Accurate prediction of protein structures and interactions using a three-track neural network. Science 2021; 373:871–876

37. Tunyasuvunakool K, Adler J, Wu Z, et al. Highly accurate protein structure prediction for the human proteome. Nature 2021; 596:590–596

38. Akdel M, Pires DEV, Porta Pardo E, et al. A structural biology community assessment of AlphaFold 2 applications. 2021;

39. He W, Wei L, Zou Q. Research progress in protein posttranslational modification site prediction. Brief. Funct. Genomics 2019; 18:220–229

40. Ding Z, Kihara D. Computational Methods for Predicting Protein-Protein Interactions Using Various Protein Features. Curr. Protoc. Protein Sci. 2018; 93:

41. Rauer C, Sen N, Waman VP, et al. Computational approaches to predict protein functional families and functional sites. Curr. Opin. Struct. Biol. 2021; 70:108–122

42. Greener JG, Sternberg MJ. Structure-based prediction of protein allostery. Curr. Opin. Struct. Biol. 2018; 50:1–8

43. Schymkowitz J, Borg J, Stricher F, et al. The FoldX web server: an online force field. Nucleic Acids Res. 2005; 33:W382–W388

44. Jespers W, Isaksen GV, Andberg TAH, et al. QresFEP: An Automated Protocol for Free Energy Calculations of Protein Mutations in Q. J. Chem. Theory Comput. 2019; 15:5461– 5473

45. Steinbrecher T, Zhu C, Wang L, et al. Predicting the Effect of Amino Acid Single-Point Mutations on Protein Stability-Large-Scale Validation of MD-Based Relative Free Energy Calculations. J. Mol. Biol. 2017; 429:948–963

46. Gapsys V, Michielssens S, Seeliger D, et al. Accurate and Rigorous Prediction of the Changes in Protein Free Energies in a Large-Scale Mutation Scan. Angew. Chem. Int. Ed Engl. 2016; 55:7364–7368

47. Pejaver V, Urresti J, Lugo-Martinez J, et al. Inferring the molecular and phenotypic impact of amino acid variants with MutPred2. Nat. Commun. 2020; 11:5918

48. Frazer J, Notin P, Dias M, et al. Disease variant prediction with deep generative models of evolutionary data. Nature 2021; 599:91–95

49. Kircher M, Witten DM, Jain P, et al. A general framework for estimating the relative pathogenicity of human genetic variants. Nat. Genet. 2014; 46:310–315

50. Almqvist J, Huang Y, Hovmöller S, et al. Homology Modeling of the Human Microsomal Glucose 6-Phosphate Transporter Explains the Mutations That Cause the Glycogen Storage Disease Type Ib. Biochemistry 2004; 43:9289–9297

51. Ittisoponpisan S, Islam SA, Khanna T, et al. Can Predicted Protein 3D Structures Provide Reliable Insights into whether Missense Variants Are Disease Associated? J. Mol. Biol. 2019; 431:2197–2212

52. Berman HM. The Protein Data Bank. Nucleic Acids Res. 2000; 28:235–242

53. Laskowski RA, Stephenson JD, Sillitoe I, et al. VarSite: Disease variants and protein structure. Protein Sci. Publ. Protein Soc. 2020; 29:111–119

54. Mi H, Muruganujan A, Casagrande JT, et al. Large-scale gene function analysis with the PANTHER classification system. Nat. Protoc. 2013; 8:1551–1566

55. Thomas PD, Ebert D, Muruganujan A, et al. PANTHER: Making genome-scale phylogenetics accessible to all. Protein Sci. Publ. Protein Soc. 2022; 31:8–22

56. Mi H, Ebert D, Muruganujan A, et al. PANTHER version 16: a revised family classification, tree-based classification tool, enhancer regions and extensive API. Nucleic Acids Res. 2021; 49:D394–D403

57. Orengo C, Michie A, Jones S, et al. CATH – a hierarchic classification of protein domain structures. Structure 1997; 5:1093–1109

58. Sillitoe I, Bordin N, Dawson N, et al. CATH: increased structural coverage of functional space. Nucleic Acids Res. 2021; 49:D266–D273

59. El-Gebali S, Mistry J, Bateman A, et al. The Pfam protein families database in 2019. Nucleic Acids Res. 2019; 47:D427–D432

60. Dessailly BH, Nair R, Jaroszewski L, et al. PSI-2: Structural Genomics to Cover Protein Domain Family Space. Structure 2009; 17:869–881

61. Das S, Lee D, Sillitoe I, et al. Functional classification of CATH superfamilies: a domain-based approach for protein function annotation. Bioinformatics 2015; 31:3460–3467

62. Medvedev KE, Kinch LN, Dustin Schaeffer R, et al. A Fifth of the Protein World: Rossmann-like Proteins as an Evolutionarily Successful Structural unit. J. Mol. Biol. 2021; 433:166788

63. Halaby DM, Poupon A, Mornon J-P. The immunoglobulin fold family: sequence analysis and 3D structure comparisons. Protein Eng. Des. Sel. 1999; 12:563–571

64. Nallapareddy V, Bordin N, Sillitoe I, et al. CATHe: Detection of remote homologues for CATH superfamilies using embeddings from protein language models. 2022;

65. Elnaggar A, Heinzinger M, Dallago C, et al. ProtTrans: Towards Cracking the Language of Life’s Code Through Self-Supervised Deep Learning and High Performance Computing. 2020;

66. Mariani V, Biasini M, Barbato A, et al. lDDT: a local superposition-free score for comparing protein structures and models using distance difference tests. Bioinformatics 2013; 29:2722–2728

67. Valdar WSJ. Scoring residue conservation. Proteins Struct. Funct. Genet. 2002; 48:227– 241

68. Mészáros B, Erdős G, Dosztányi Z. IUPred2A: context-dependent prediction of protein disorder as a function of redox state and protein binding. Nucleic Acids Res. 2018; 46:W329–W337

69. Schriml LM, Munro JB, Schor M, et al. The Human Disease Ontology 2022 update. Nucleic Acids Res. 2022; 50:D1255–D1261

70. Sevim Bayrak C, Stein D, Jain A, et al. Identification of discriminative gene-level and protein-level features associated with pathogenic gain-of-function and loss-of-function variants. Am. J. Hum. Genet. 2021; 108:2301–2318

71. Stenson PD, Mort M, Ball EV, et al. The Human Gene Mutation Database (HGMD®): optimizing its use in a clinical diagnostic or research setting. Hum. Genet. 2020; 139:1197– 1207

72. Esposito D, Weile J, Shendure J, et al. MaveDB: an open-source platform to distribute and interpret data from multiplexed assays of variant effect. Genome Biol. 2019; 20:223

73. Campbell AJ, Watts KJ, Johnson MS, et al. Gain-of-function mutations cluster in distinct regions associated with the signalling pathway in the PAS domain of the aerotaxis receptor, Aer: Signalling in the Aer-PAS domain. Mol. Microbiol. 2010; 77:575–586

74. Kamburov A, Lawrence MS, Polak P, et al. Comprehensive assessment of cancer missense mutation clustering in protein structures. Proc. Natl. Acad. Sci. 2015; 112:E5486– E5495

75. Meyer MJ, Lapcevic R, Romero AE, et al. mutation3D: Cancer Gene Prediction Through Atomic Clustering of Coding Variants in the Structural Proteome. Hum. Mutat. 2016; 37:447– 456

76. Vacic V, Uversky VN, Dunker AK, et al. Composition Profiler: a tool for discovery and visualization of amino acid composition differences. BMC Bioinformatics 2007; 8:211

77. Gao M, Zhou H, Skolnick J. Insights into Disease-Associated Mutations in the Human Proteome through Protein Structural Analysis. Structure 2015; 23:1362–1369

78. Yang J, Kwon S, Bae S-H, et al. GalaxySagittarius: Structure- and Similarity-Based Prediction of Protein Targets for Druglike Compounds. J. Chem. Inf. Model. 2020; 60:3246– 3254

79. Singh N, Decroly E, Khatib A-M, et al. Structure-based drug repositioning over the human TMPRSS2 protease domain: search for chemical probes able to repress SARS-CoV-2 Spike protein cleavages. Eur. J. Pharm. Sci. 2020; 153:105495

80. Xue LC, Dobbs D, Bonvin AMJJ et al. Computational prediction of protein interfaces: A review of data driven methods. FEBS Lett. 2015; 589:3516–3526

81. Lo Gullo G, De Santis ML, Paiardini A, et al. The Archaeal Elongation Factor EF-2 Induces the Release of aIF6 From 50S Ribosomal Subunit. Front. Microbiol. 2021; 12:631297

82. Diesterbeck US, Gittis AG, Garboczi DN, et al. The 2.1 Å structure of protein F9 and its comparison to L1, two components of the conserved poxvirus entry-fusion complex. Sci. Rep. 2018; 8:16807

83. Prabantu VM, Naveenkumar N, Srinivasan N. Influence of Disease-Causing Mutations on Protein Structural Networks. Front. Mol. Biosci. 2021; 7:620554

84. Chakrabarty B, Parekh N. NAPS: Network Analysis of Protein Structures. Nucleic Acids Res. 2016; 44:W375–W382

85. Jack BR, Meyer AG, Echave J, et al. Functional Sites Induce Long-Range Evolutionary Constraints in Enzymes. PLOS Biol. 2016; 14:e1002452

86. Rodrigues CHM, Pires DEV, Ascher DB. DynaMut2: Assessing changes in stability and flexibility upon single and multiple point missense mutations. Protein Sci. Publ. Protein Soc. 2021; 30:60–69

87. Savojardo C, Manfredi M, Martelli PL, et al. Solvent Accessibility of Residues Undergoing Pathogenic Variations in Humans: From Protein Structures to Protein Sequences. Front. Mol. Biosci. 2021; 7:626363

88. Iqbal S, Pérez-Palma E, Jespersen JB, et al. Comprehensive characterization of amino acid positions in protein structures reveals molecular effect of missense variants. Proc. Natl. Acad. Sci. 2020; 117:28201–28211

89. Mann HB, Whitney DR. On a Test of Whether one of Two Random Variables is Stochastically Larger than the Other. Ann. Math. Stat. 1947; 18:50–60

90. Porta-Pardo E, Ruiz-Serra V, Valentini S, et al. The structural coverage of the human proteome before and after AlphaFold. PLOS Comput. Biol. 2022; 18:e1009818

91. Sen N, Kanitkar TR, Roy AA, et al. Predicting and designing therapeutics against the Nipah virus. PLoS Negl. Trop. Dis. 2019; 13:e0007419

92. Nguyen MN, Sen N, Lin M, et al. Discovering Putative Protein Targets of Small Molecules: A Study of the p53 Activator Nutlin. J. Chem. Inf. Model. 2019; 59:1529–1546

93. Waman VP, Sen N, Varadi M, et al. The impact of structural bioinformatics tools and resources on SARS-CoV-2 research and therapeutic strategies. Brief. Bioinform. 2021; 22:742–768

94. Farheen N, Sen N, Nair S, et al. Depth dependent amino acid substitution matrices and their use in predicting deleterious mutations. Prog. Biophys. Mol. Biol. 2017; 128:14–23

95. Kanitkar TR, Sen N, Nair S, et al. Methods for Molecular Modelling of Protein Complexes. Struct. Proteomics 2021; 2305:53–80

96. Ashford P, Pang CSM, Moya-García AA, et al. A CATH domain functional family based approach to identify putative cancer driver genes and driver mutations. Sci. Rep. 2019; 9:263

97. Das S, Scholes HM, Sen N, et al. CATH functional families predict functional sites in proteins. Bioinformatics 2021; 37:1099–1106

98. Altschul SF, Gish W, Miller W, et al. Basic local alignment search tool. J. Mol. Biol. 1990; 215:403–410

99. Mistry J, Finn RD, Eddy SR, et al. Challenges in homology search: HMMER3 and convergent evolution of coiled-coil regions. Nucleic Acids Res. 2013; 41:e121–e121

100. Lewis TE, Sillitoe I, Lees JG. cath-resolve-hits: a new tool that resolves domain matches suspiciously quickly. Bioinformatics 2019; 35:1766–1767

101. Katoh K, Standley DM. MAFFT Multiple Sequence Alignment Software Version 7: Improvements in Performance and Usability. Mol. Biol. Evol. 2013; 30:772–780

102. Steinegger M, Meier M, Mirdita M, et al. HH-suite3 for fast remote homology detection and deep protein annotation. BMC Bioinformatics 2019; 20:473

103. Mirdita M, von den Driesch L, Galiez C, et al. Uniclust databases of clustered and deeply annotated protein sequences and alignments. Nucleic Acids Res. 2017; 45:D170– D176

104. Steinegger M, Mirdita M, Söding J. Protein-level assembly increases protein sequence recovery from metagenomic samples manyfold. Nat. Methods 2019; 16:603–606

105. Steinegger M, Söding J. Clustering huge protein sequence sets in linear time. Nat. Commun. 2018; 9:2542

106. Anishchenko I, Baek M, Park H, et al. Protein tertiary structure prediction and refinement using deep learning and Rosetta in CASP14. Proteins Struct. Funct. Bioinforma. 2021; prot.26194

107. Chaudhury S, Lyskov S, Gray JJ. PyRosetta: a script-based interface for implementing molecular modeling algorithms using Rosetta. Bioinformatics 2010; 26:689–691

108. Hiranuma N, Park H, Baek M, et al. Improved protein structure refinement guided by deep learning based accuracy estimation. Nat. Commun. 2021; 12:1340

109. Zemla A. LGA: a method for finding 3D similarities in protein structures. Nucleic Acids Res. 2003; 31:3370–3374

110. Park H, Bradley P, Greisen P, et al. Simultaneous Optimization of Biomolecular Energy Functions on Features from Small Molecules and Macromolecules. J. Chem. Theory Comput. 2016; 12:6201–6212

111. Krissinel E. Enhanced fold recognition using efficient short fragment clustering. J. Mol. Biochem. 2012; 1:76–85

112. Kwon S, Won J, Kryshtafovych A, et al. Assessment of protein model structure accuracy estimation in CASP14 : Old and new challenges. Proteins Struct. Funct. Bioinforma. 2021; 89:1940–1948

113. Soni N. neeleshsoni21/Cell_list. 2021;

114. Yao Z, Wang J-S, Liu G-R, et al. Improved neighbor list algorithm in molecular simulations using cell decomposition and data sorting method. Comput. Phys. Commun. 2004; 161:27–35

115. Dobson M, Fox I, Saracino A. Cell List Algorithms for Nonequilibrium Molecular Dynamics. ArXiv14123784 Phys. 2014;

116. Dhawanjewar AS, Roy AA, Madhusudhan MS. A knowledge-based scoring function to assess quaternary associations of proteins. Bioinformatics 2020; 36:3739–3748

117. Krivák R, Hoksza D. P2Rank: machine learning based tool for rapid and accurate prediction of ligand binding sites from protein structure. J. Cheminformatics 2018; 10:39

118. Qin S, Zhou H-X. meta-PPISP: a meta web server for protein-protein interaction site prediction. Bioinformatics 2007; 23:3386–3387

119. Naccess homepage.

120. Heberle H, Meirelles GV, da Silva FR, et al. InteractiVenn: a web-based tool for the analysis of sets through Venn diagrams. BMC Bioinformatics 2015; 16:169

