## Supplementary Data for "Characterizing and explaining impact of disease-associated mutations in proteins without known structures or structural homologues"

### Supplementary Information

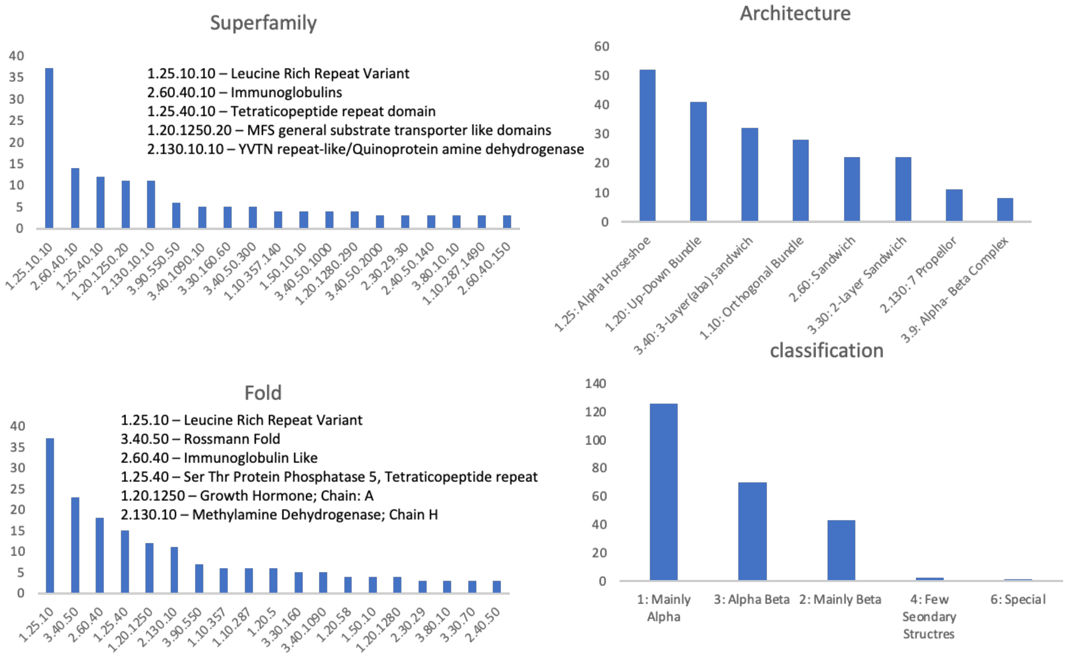


Figure S1 - Frequency of modelled CATH domains across the CATH hierarchy.

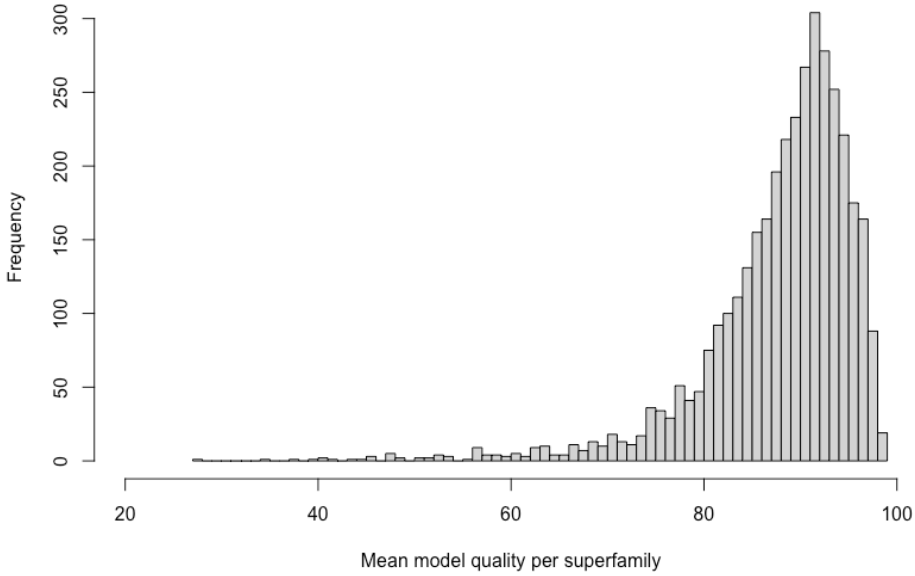


Figure S2- Mean model quality of the CATH superfamilies for the AlphaFold models.

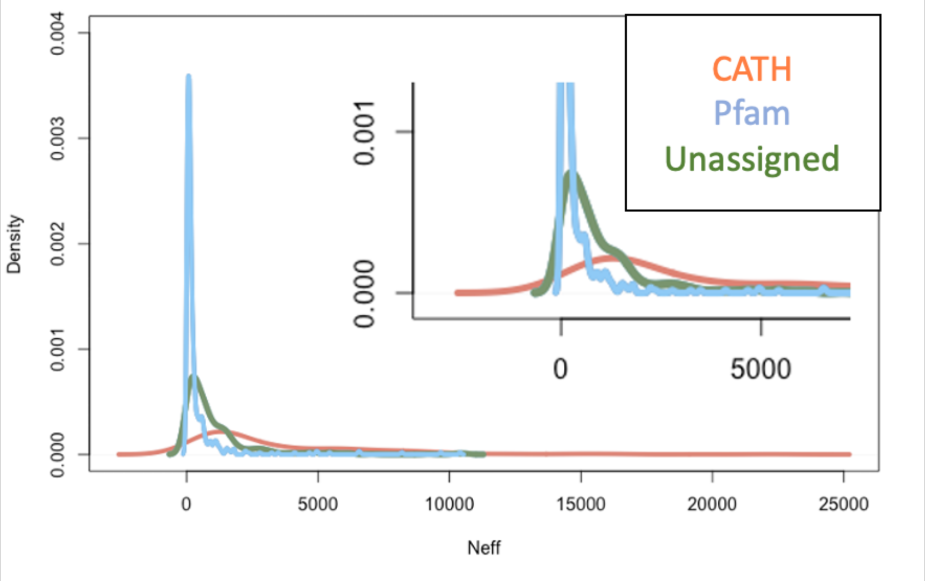


Figure S3- Density plot showing the Neff values of the MSA for CATH domains in red, Pfam domains in green and unassigned domains in blue. The plot within is a zoomed version for the neff range 0-5000 and density range 0-0.001.

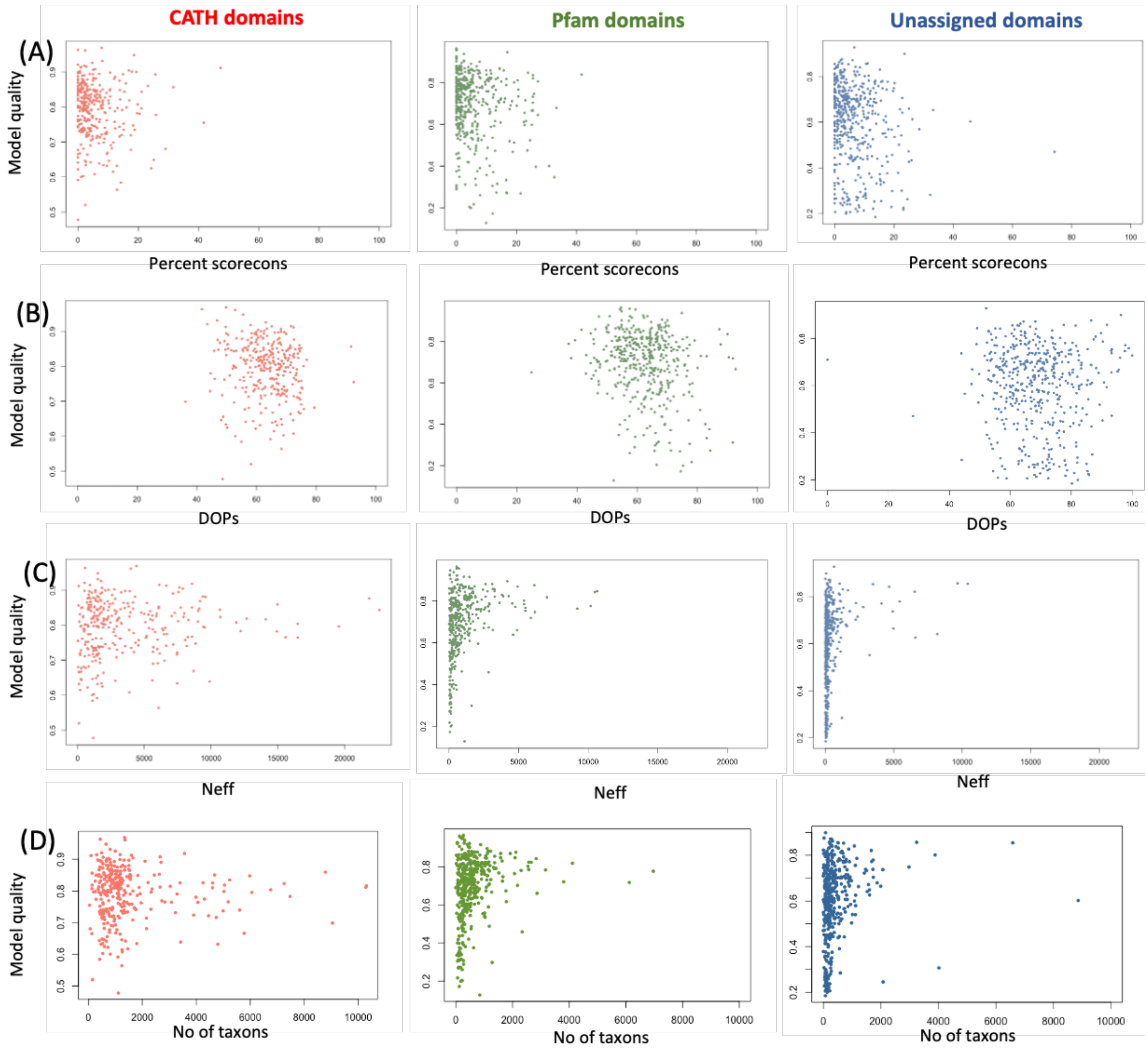


Figure S4 - Plot showing the different alignment quality parameters (A) Percent Scorecons (B) DOPs (C) Neff (D) No of Taxon ID in the alignment vs the RoseTTAFold model quality for CATH domains (in red), Pfam domains (in green) and unassigned domains (in blue).

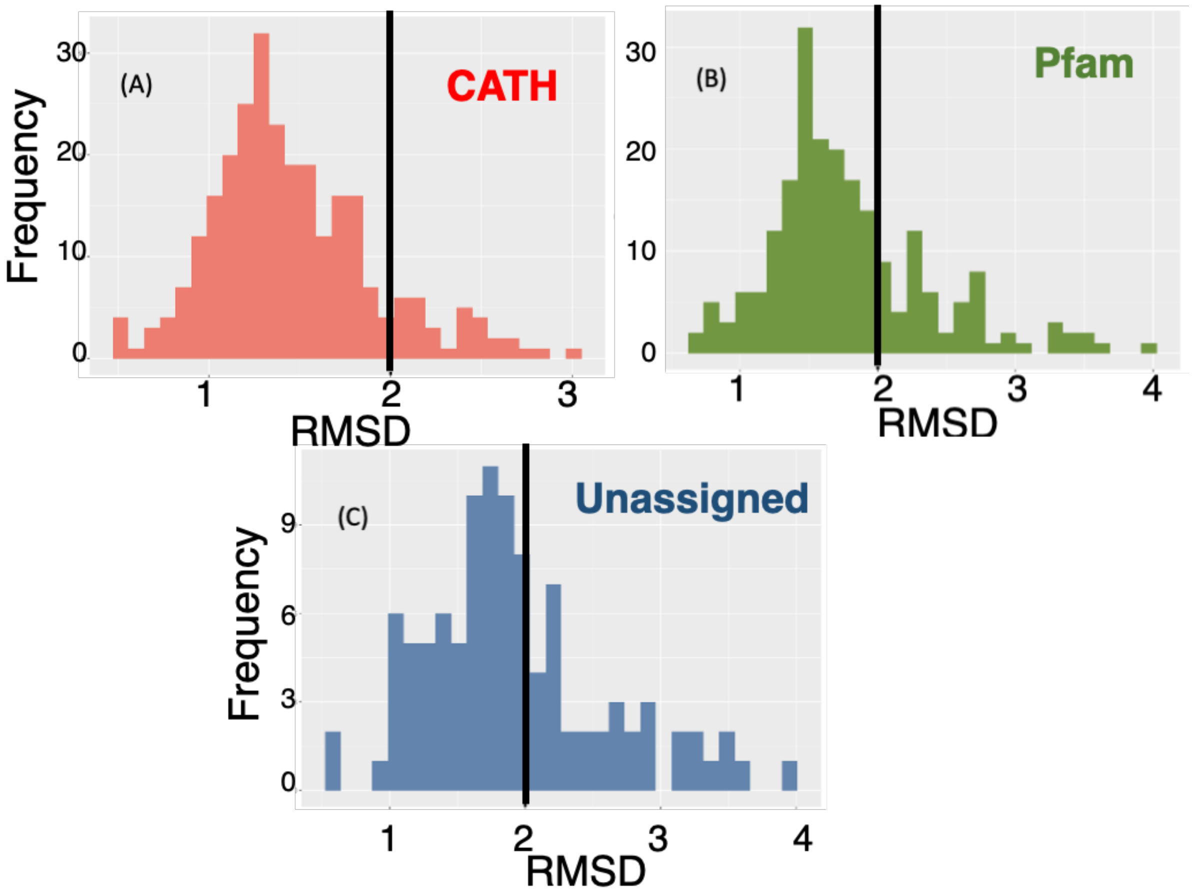


Figure S5 - RMSD (Å) between AlphaFold and RoseTTAFold models for A) CATH domains B) Pfam domains and C) Unassigned domains. The dotted line indicates 2 Å RMSD.

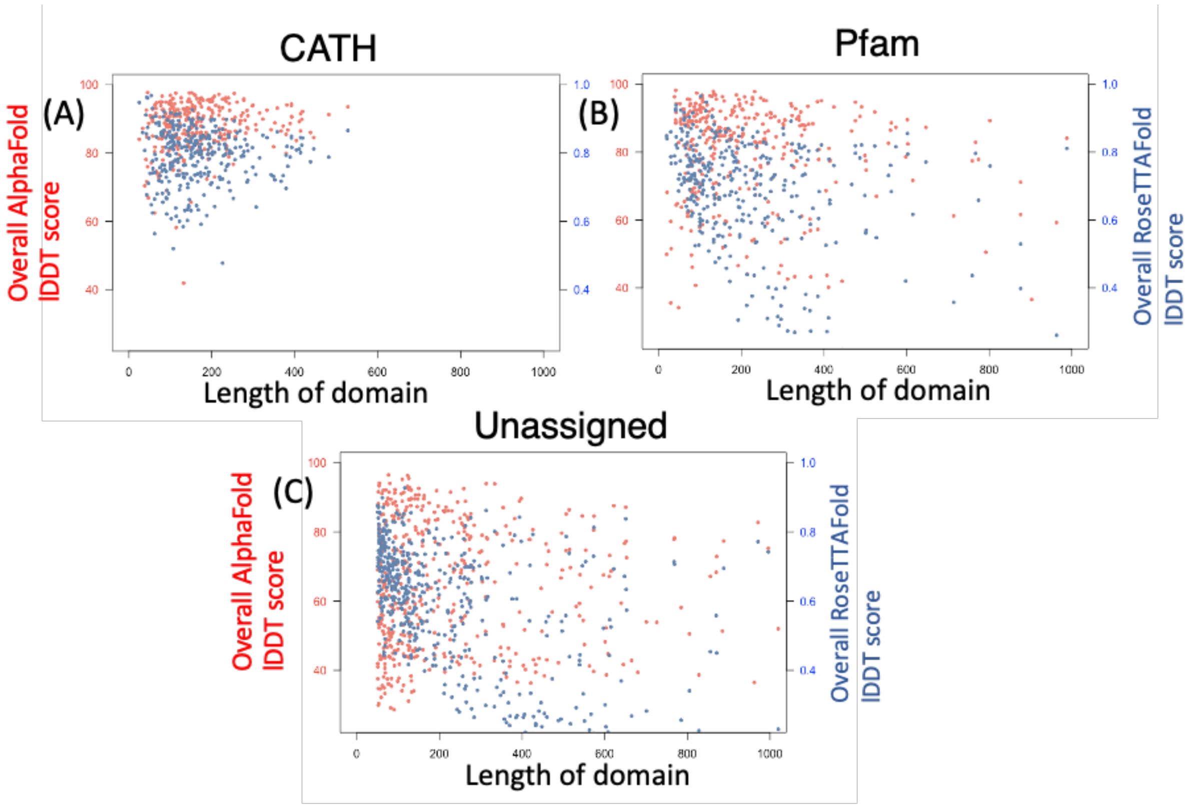


Figure S6 - Distribution of model quality (AlphaFold in red, RoseTTAFold in blue) versus the length of the modelled domain along the x-axis for a) CATH domains b) Pfam domains and c) Unassigned domains

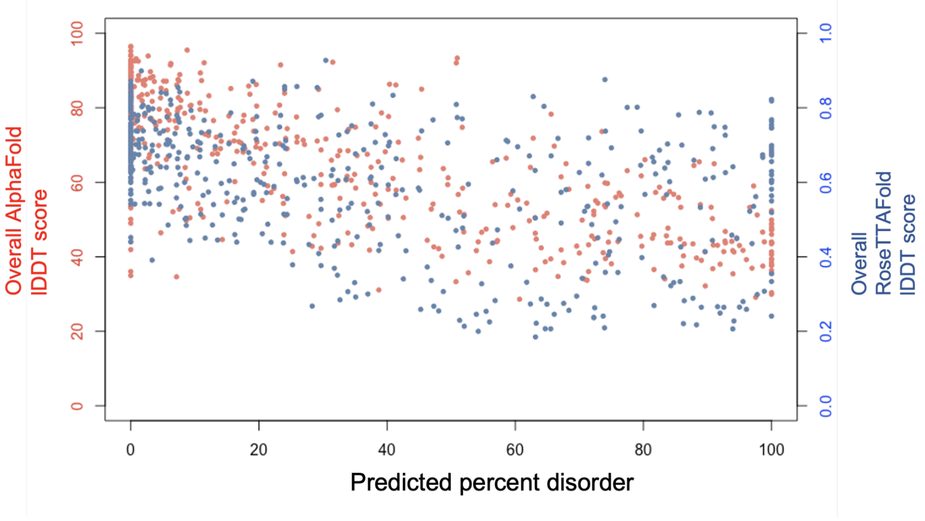


Figure S7 - Distribution of model quality (AlphaFold in red, RoseTTAFold in blue) against the predicted percent disorder along the x-axis for Unassigned domains.

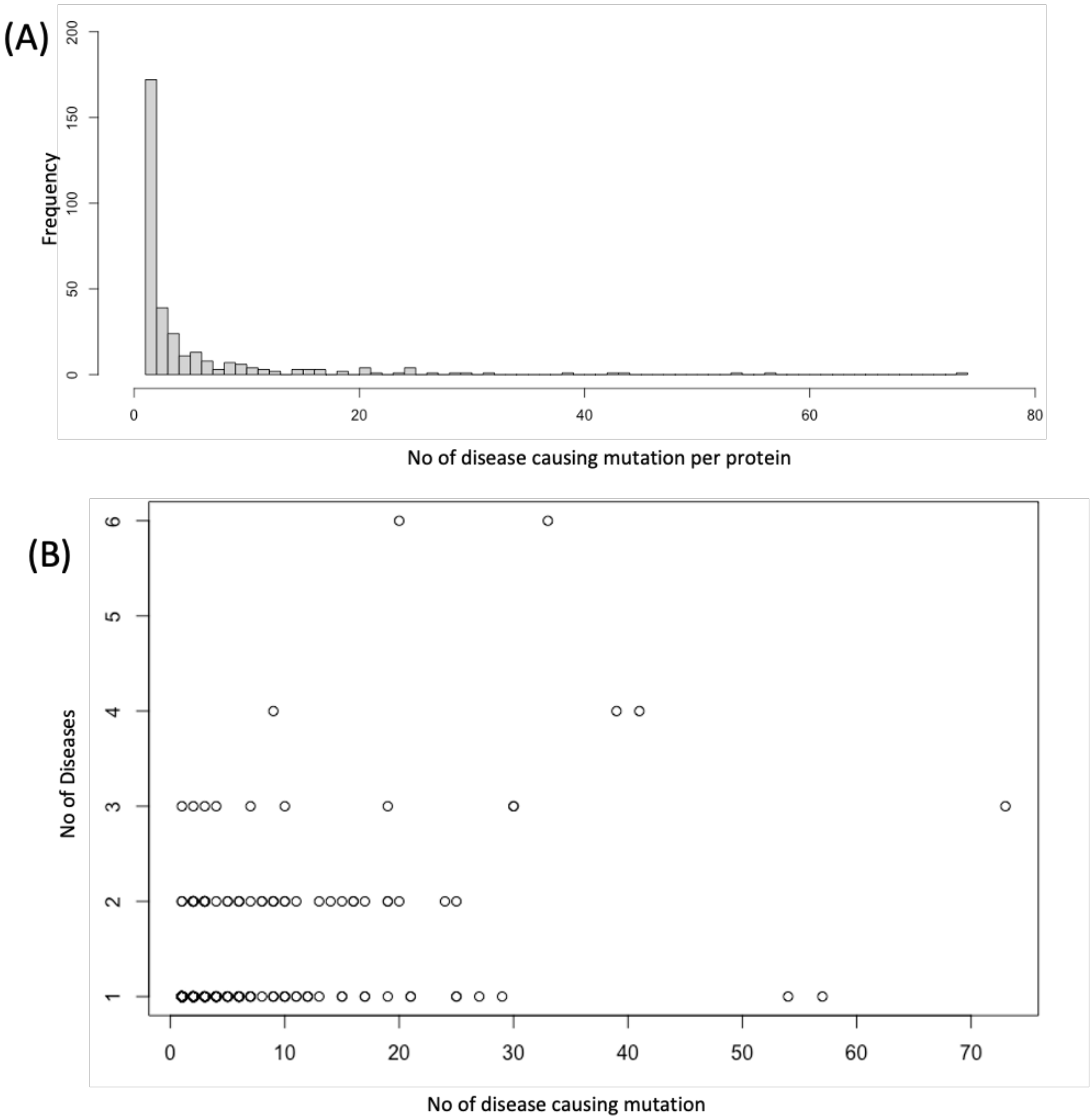


Figure S8 - (A) Number of disease-associated mutations per protein in humsavar. (B) Number of unique mutations per protein vs the number of unique diseases per protein

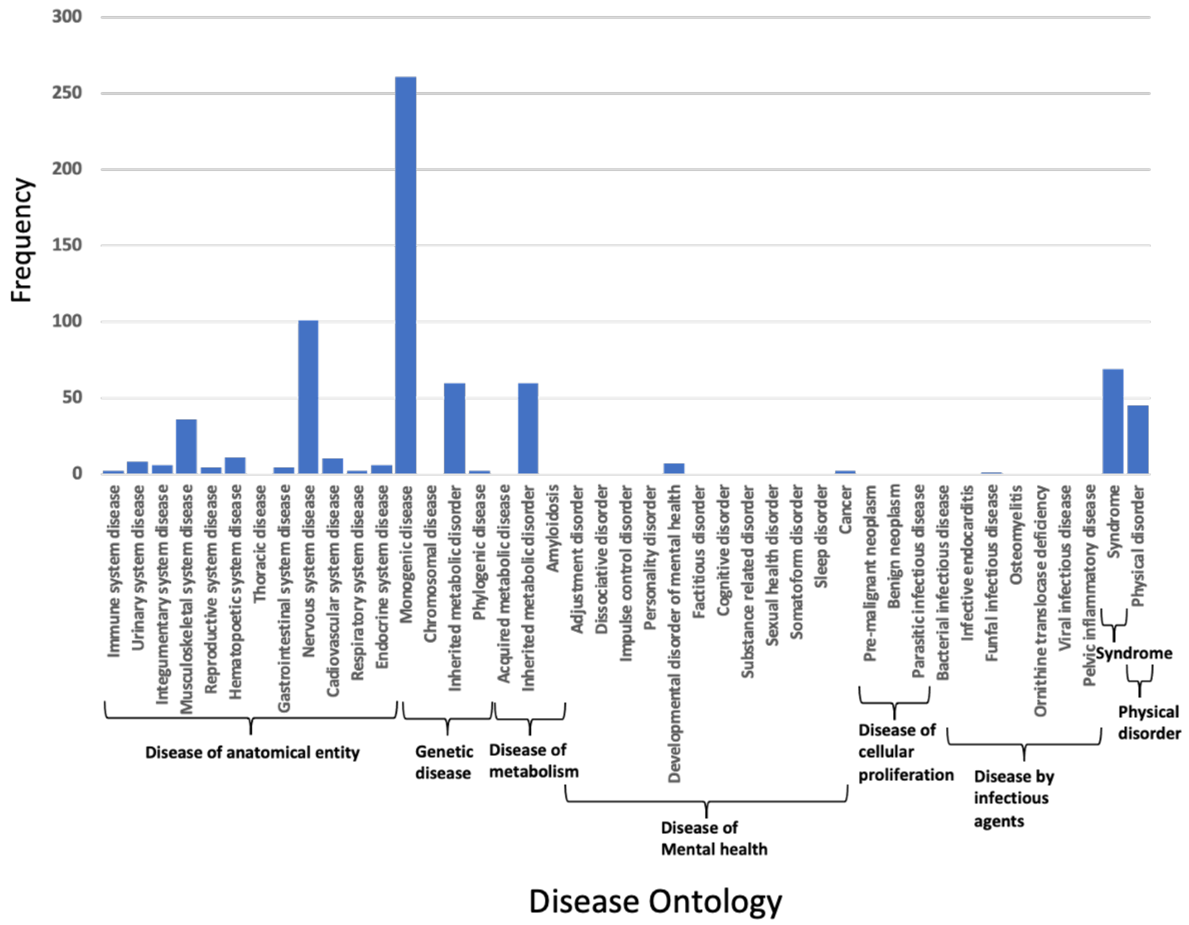


Figure S9 - Frequency of the various diseases in the study dataset belonging to the various disease ontologies. The disease ontologies and their groupings can be found here.

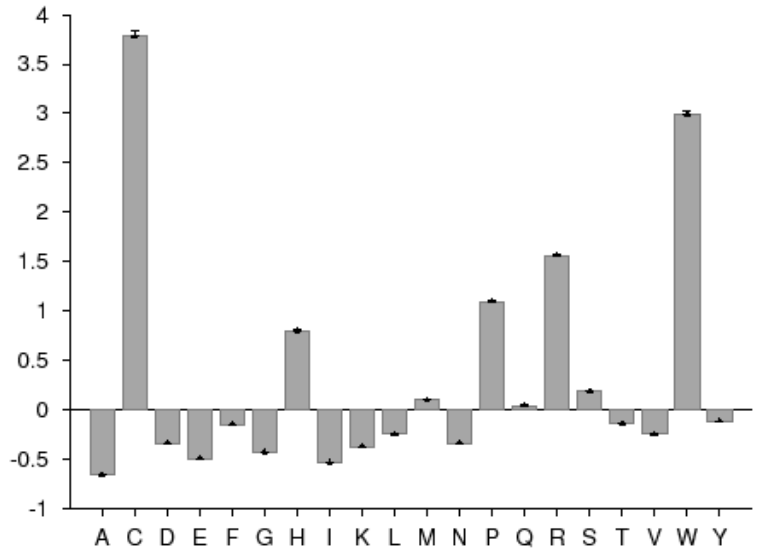


Figure S10 - Relative abundance of the mutant amino acids that are disease associated.

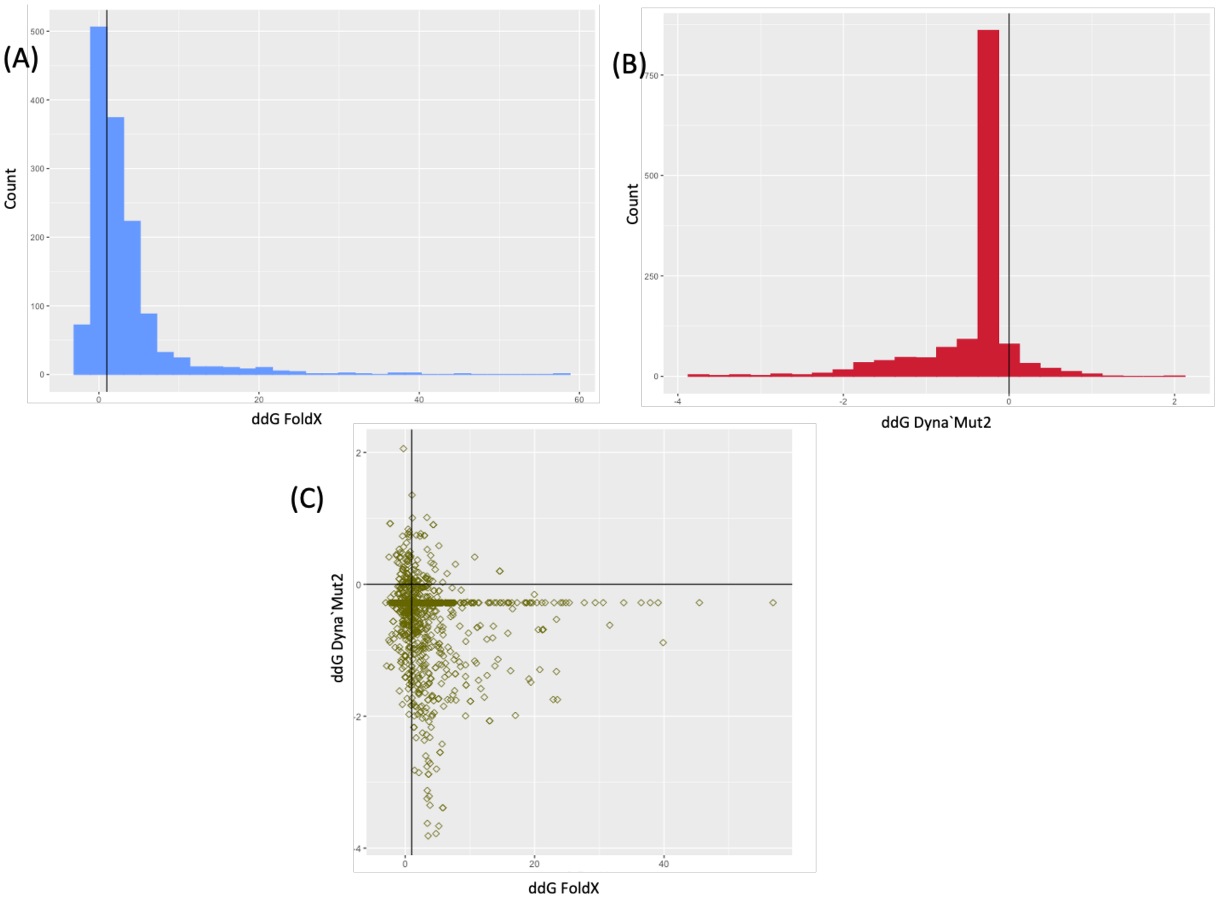


Figure S11 - Histogram of ddG of the disease-associated mutations calculated from AlphaFold domain models calculated using A) FoldX B) Dynamut2. The black vertical line indicates ddG=1 and ddG=0 respectively (ddG>1 was chosen as destabilising as per FoldX because FoldX has a larger range of ddG and a positive ddG indicates a destabilising mutation; for DynaMut2 ddG<0 indicates destabilising mutation because it has a much smaller range and negative ddG indicates destabilising mutations). C) Scatter plot showing the correlatin between the FoldX and Dynamut2 predictions.

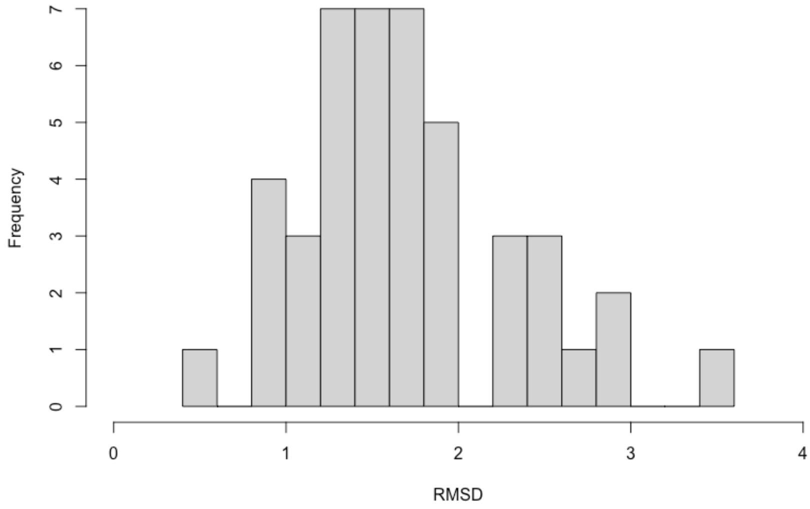


Figure S12 - Histogram of RMSD between AlphaFold and RoseTTAFold models for the 45 proteins which had 93 mutations which could be explained using RosettaFold but not AlphaFold

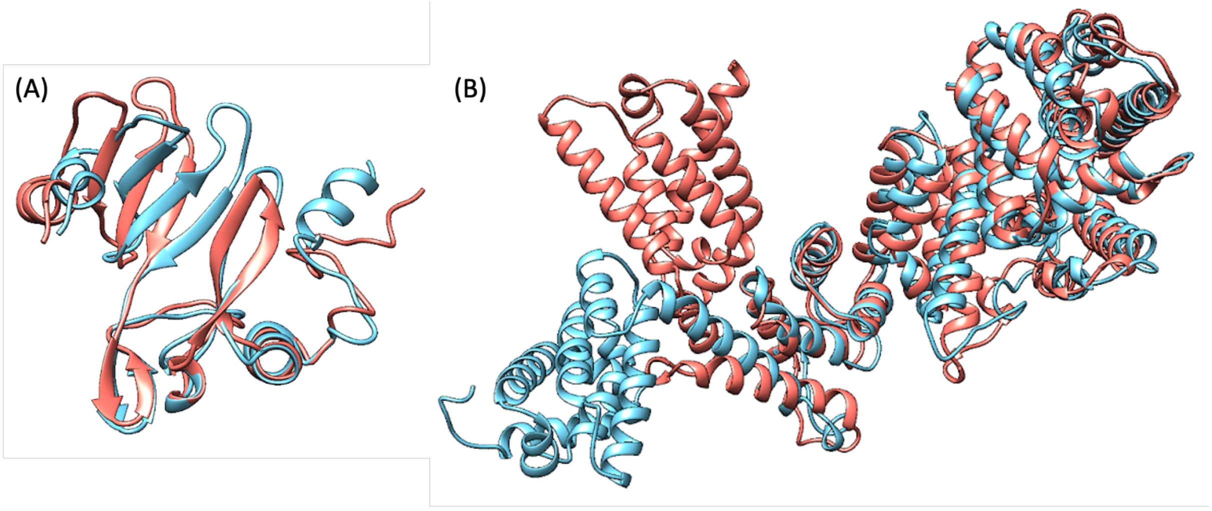


Figure S13 - Structural superimposition of AlphaFold model in salmon and RoseTTAFold model in blue for a) Fukutin-related protein (Uniprot ID Q9H9S5) (residue nos 374-495) b) Fanconi Anaemia Group C protein (Uniprot ID Q00597) assigned to PFAM FunFam PF02106.15_Fanconi_C-FF-000001

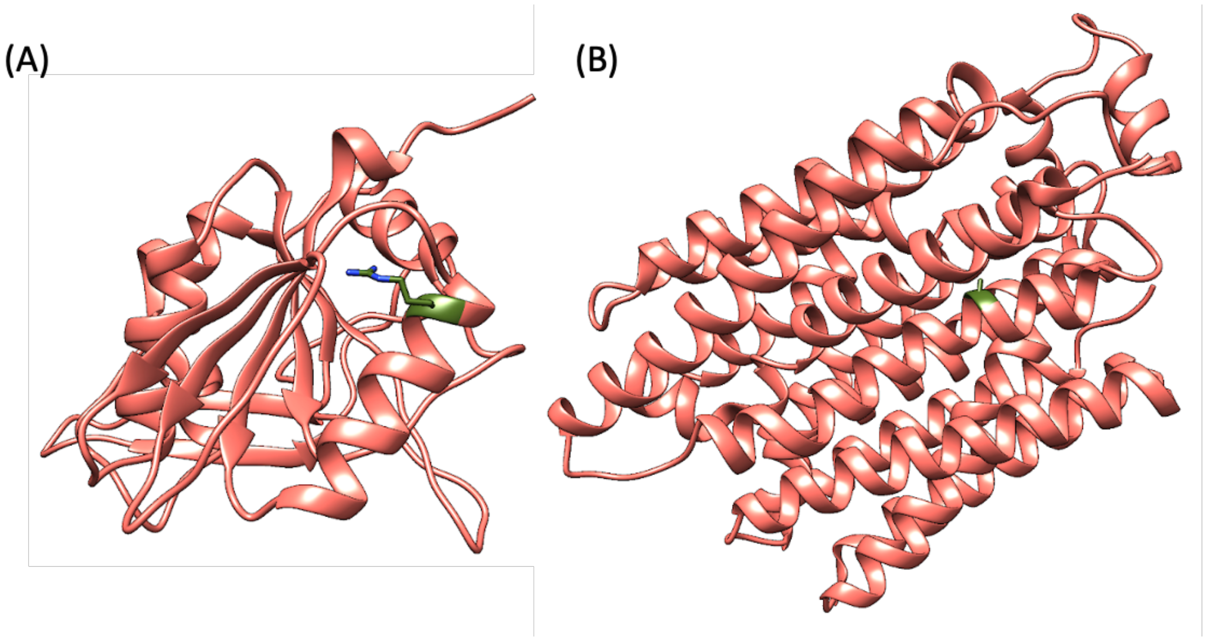


Figure S14 - AlphaFold model in salmon of a) Beta-1,4 N-acetylgalactosaminyltransferase 1 (Q00973) assigned to CATH FunFam 3.90.550.10-FF-000076, where mutation Arg300Cys (colored in olive green) leads to a disease b) Feline leukaemia virus subgroup C receptor-related protein 1(Q9Y5Y0) mapped to CATH FunFam 1.20.1250.20-FF-000184 where mutation of Ala241Thr (colored in olive green) leads to a disease

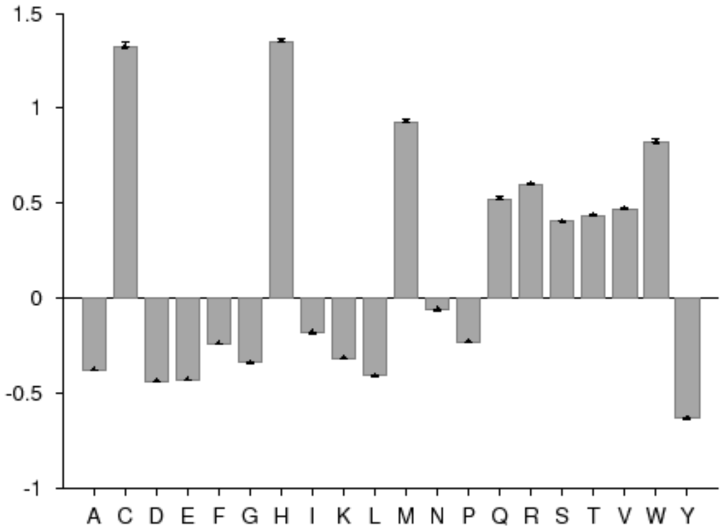


Figure S15 - Relative abundance of the mutant amino acids associated with polymorphism.

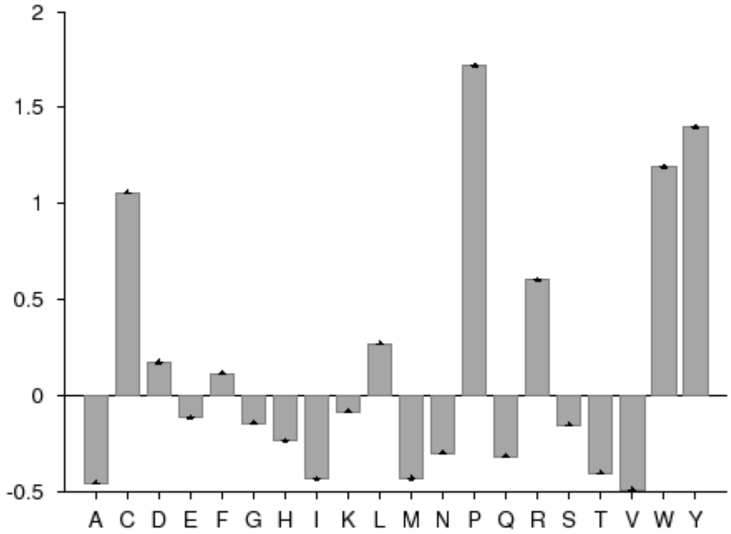


Figure S16 - Relative abundance of amino acids of the deleterious mutations with the residues in polymorphism as the background.

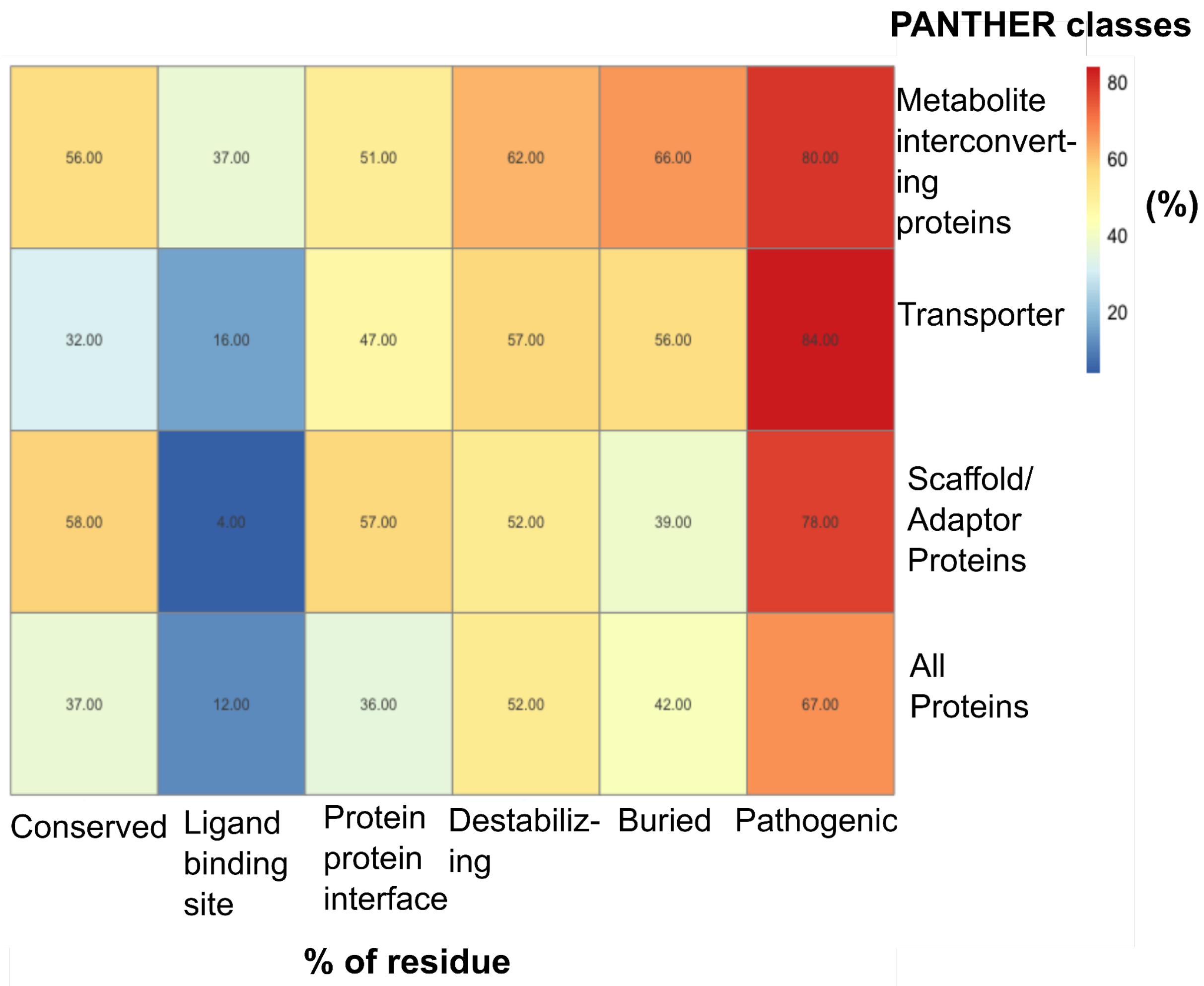


Figure S17 - Heatmap showing the percentage of disease associated residues that are near predicted conserved, ligand binding site, protein-protein interface residues. Also shown are the percentages of destabilising, buried and pathogenic disease associated mutations for 3 PANTHER classes and across all proteins.

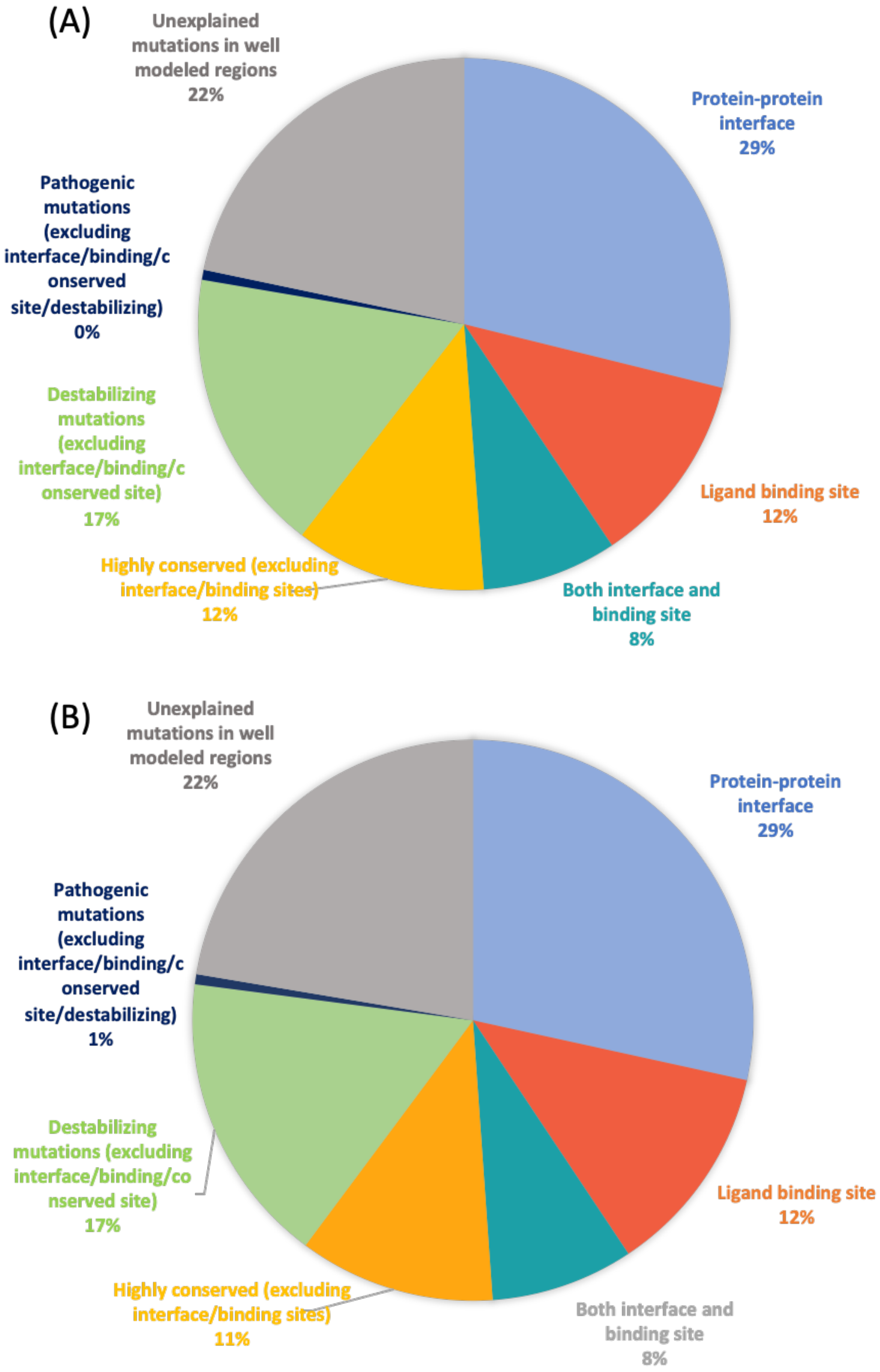


Figure S18 - Piechart showing the percentage of mutations in the well modelled regions that can be explained based on proximity to functional sites/destabilisation/pathogenicity for (A) dataset used in our study (B) excluding domains where MODELLER identifies a homologue in the PDB (with at least 50% sequence coverage and 30% sequence identity)

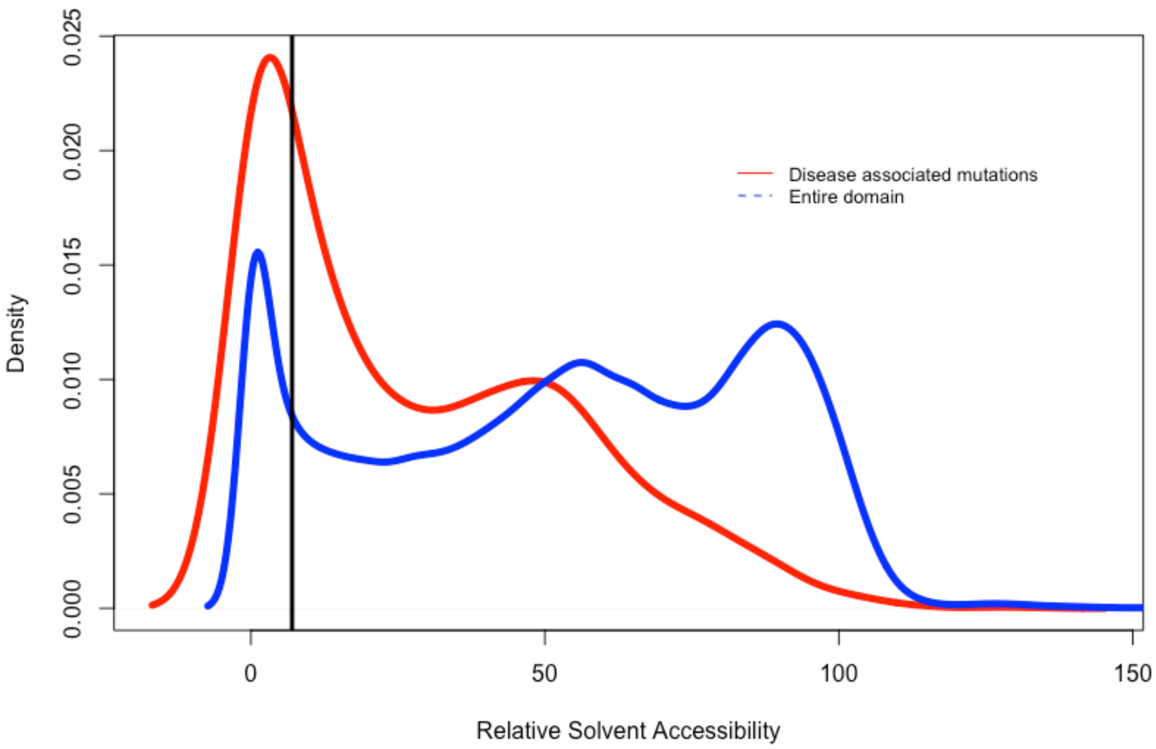


Figure S19 - Density plot of the relative solvent accessibility of the residues that are disease associated in red and for the entire domain in blue.

Table S1 - Table showing the amino acids enriched and depleted in the mutants

| | | |
|---|---|---|
| A | Depleted. | P-value=0.000000 (≤0.050000) |
| C | Enriched. | P-value=0.000000 (≤0.050000) |
| D | Depleted. | P-value=0.000728 (≤0.050000) |
| E | Depleted. | P-value=0.000000 (≤0.050000) |
| F | Not significant. | P-value=0.192748 (>0.050000) |
| G | Depleted. | P-value=0.000001 (≤0.050000) |
| H | Enriched. | P-value=0.000000 (≤0.050000) |
| I | Depleted. | P-value=0.000000 (≤0.050000) |
| K | Depleted. | P-value=0.000088 (≤0.050000) |
| L | Depleted. | P-value=0.000612 (≤0.050000) |
| M | Not significant. | P-value=0.550265 (>0.050000) |
| N | Depleted. | P-value=0.003071 (≤0.050000) |
| P | Enriched. | P-value=0.000000 (≤0.050000) |
| Q | Not significant. | P-value=0.748053 (>0.050000) |
| R | Enriched. | P-value=0.000000 (≤0.050000) |
| S | Enriched. | P-value=0.038397 (≤0.050000) |

| | | |
|---|---|---|
| T | Not significant. | P-value=0.146221 (>0.050000) |
| V | Depleted. | P-value=0.004612 (≤0.050000) |
| W | Enriched. | P-value=0.000000 (≤0.050000) |
| Y | Not significant. | P-value=0.364642 (>0.050000) |

Table S2 - Panther protein classes and their IDs and the proteins in each class

| PANTHER Class name | PANTHER Class ID | Uniprot ID |
|---|---|---|
| DNA metabolism protein | PC00009 | P52926,Q8N6T0 |
| RNA metabolism protein | PC00031 | O75448,P05549,Q12789,Q17RN3,Q2VPK5,Q53R41,Q7Z6J9,Q92481,Q99700,Q9NVC6,Q9NYY8,Q9ULK4 |
| calcium-binding protein | PC00060 | |
| cell adhesion molecule | PC00069 | O43556,Q07507,Q16586,Q7RTU9 |
| cell junction protein | PC00070 | |
| chaperone | PC00072 | A6NFY7,O00623,Q15526,Q5U5X0,Q6UW78,Q7KZN9,Q7Z412,Q86WW8,Q86Y56,Q8N9W5,Q9BRT2,Q9BTW9,Q9Y244,Q9Y2R0 |
| chromatin/chromatin-binding, or - regulatory protein | PC00077 | P42695,Q1MSJ5,Q9BV73,Q9BZW7,Q9P267 |
| cytoskeletal protein | PC00085 | O95995,Q13326,Q16585,Q5TD94,Q702N8,Q7Z3E5,Q8IYY4,Q92629,Q9NPC6,Q9UPV0 |
| defense/immunity protein | PC00090 | A6NNB3,P11912,Q86SU0 |
| extracellular matrix protein | PC00102 | Q17R60,Q32P28,Q8IVL5 |
| gene-specific transcriptional regulator | PC00264 | O00712,Q12857,Q14938,Q2TAL8,Q5JUK2,Q6ZN30,Q8NI27,Q8WUU5,Q96D09,Q9H869 |
| intercellular signal molecule | PC00207 | P01148,P01213,Q14956 |
| membrane traffic protein | PC00150 | O14653,P08247,Q12768,Q13286,Q5VIR6,Q8NEZ2,Q92734,Q96A65,Q96HR9,Q9BRK0,Q9H902,Q9UL33,Q9UPV9 |
| metabolite interconversion protein | PC00262 | A6NFK2,A8MXD5,B0YJ81,O15121,O15228,O43505,O75845,O75907,P27544,P28300,P31213,P35575,P48651,P53701,P57054,Q08397,Q12887,Q14442,Q15800,Q16635,Q1HG44,Q3MUY2,Q49MI3,Q5NDL2,Q5RI15,Q5VTY9,Q6P2C0,Q6ZMB0,Q7LGC8,Q7Z7B1,Q86WG3,Q86X52,Q8IV08,Q8N8W4,Q8NAT1,Q8NCE2,Q8NCH0,Q8NCR0,Q8NEL9,Q92535,Q92685,Q96AD5,Q96CG8,Q96EU7,Q96F25,Q96L58,Q96N66,Q9BQB6,Q9BRB3,Q9BT22,Q9BUM1,Q9BV10,Q9BVK2,Q9BZ71,Q9GZR5,Q9GZX3,Q9H237,Q9H2A9,Q9H3S5,Q9H6U8,Q9HAT2,Q9NPF2,Q9NST1,Q9NYP7,Q9P2X0,Q9UBY8,Q9Y2B1,Q9Y2B2,Q9Y4D2,Q9Y5Z9,Q9Y672 |
| protein modifying enzyme | PC00260 | O43462,O75718,P60484,P56180,Q6XPS3,Q13064,Q6XUX3,Q8NBJ5,Q8WVT3,Q96T52,Q9NUD9,Q9UHJ9,Q9Y4F9 |

| | | |
|---|---|---|
| protein-binding activity modulator | PC00095 | O96020,P24864,P46019,P46020,Q15042,Q8WYR1,Q93100,Q9H2M9,Q9UJJ9 |
| scaffold/adaptor protein | PC00226 | O43734,O95859,Q9UQC2,Q13480,P23942,P48509,P56539,Q03135,Q03395,Q08AM6,Q6UX65,Q7Z6L0,Q8WWZ3,Q96FG2,Q99732,Q9C004 |
| storage protein | PC00210 | |
| structural protein | PC00211 | A5D8W1,P60201,Q86UC2,Q96M91,Q96RY7,Q9BPU9,Q9BRQ4,Q9NRM1,Q9NWB7,Q9NXB0,Q9UPM9 |
| transfer/carrier protein | PC00219 | O14791 |
| translational protein | PC00263 | Q9H1X1 |
| transmembrane signal receptor | PC00197 | O75787,Q6PRD1,Q8WVP7,Q9BUN5,Q9GZN0 |
| transporter | PC00227 | A6NNN8,O00400,O60779,O60931,O95436,P39210,Q06495,Q08357,Q0D2K0,Q0ZLH3,Q15904,Q495M3,Q53GD3,Q6P4A7,Q75T13,Q7RTP0,Q7Z403,Q8IU68,Q8N130,Q8TDI8,Q92959,Q96A29,Q96H72,Q99567,Q9BRI3,Q9BZV2,Q9C0K1,Q9HAB3,Q9NQ11,Q9NQ40,Q9NUN5,Q9NWF4,Q9UMX9,Q9Y2W3,Q9Y5Y0,Q9Y6L6 |
| viral or transposable element protein | PC00237 | |

Table S3 - Table showing the number of mutations in the well modelled regions that can be explained based on proximity to functional sites/destabilisation/pathogenicity for - dataset used in our study and excluding domains where MODELLER identifies a homologue in the PDB (with at least 50% sequence coverage an 30% sequence identity)

| **Functional site/destabilising/pathogenic/unexplained** | **Dataset used in the study** | **Dataset after removal of proteins with homologue detected by MODELLER** |
|---|---|---|
| Protein-protein interface | 380 | 342 |
| Ligand binding site | 155 | 147 |
| Both interface and binding site | 108 | 99 |
| Highly conserved (excluding interface/binding sites) | 153 | 136 |
| Destabilising mutations (excluding interface/binding/conserved site) | 227 | 203 |
| Pathogenic mutations (excluding interface/binding/conserved site/destabilising) | 8 | 7 |
| Unexplained mutations in well modelled regions | 286 | 269 |

### Text S1 - Gene Ontology analysis

Gene ontology analysis (molecular function) using PANTHER [1] for these 553 proteins showed enrichment of some gene functions, including phosphatidylinositol N-acetylglucosaminyltransferase activity (30 fold enrichment), melanocortin receptor binding (type 1,3 and 4) (28 fold enrichment), sodium-phosphate symporter activity (13 fold enrichment). The biological processes enriched were sodium-dependent phosphate transport (37 fold enrichment), protein localisation to the ciliary transition zone (29 fold enrichment), Endoplasmic reticulum tubular network organisation (28 fold enrichment), lysosomal lumen acidification (27 fold enrichment), attachment of GPI anchor to proteins (25 fold enrichment). The GO enrichment can be explained because a large number of these proteins are either metabolite interconverting enzymes, transporter or scaffold/adaptor proteins. The enrichment in transport proteins could therefore be explained by the fact that transport proteins are mainly transmembrane and difficult to crystallise.

### Text S2 - MSA diversity does not correlate with model quality

AlphaFold reports that with an increase in the number of sequences beyond 100 there is a threshold effect which only leads to a small improvement for increasing numbers of sequences [2]**.** RoseTTAFold authors have reported similar trends [3]. The authors of these techniques hypothesise that the MSA information is useful in the early stages of the network to get a coarse overview of the structure, however, the final refinement does not depend on the MSA diversity.

### Text S3 - Characterising disease-associated mutations

Around 81%, 14% and 3% of these proteins have only 1, 2 and 3 disease annotations each. Only 3 and 2 proteins had mutations leading to 4 and 6 different diseases each respectively. 40 of the proteins had more than 10 disease-associated mutations (Figure S8A). Of these 19, 13, 4, 2 and 2 proteins had 1,2,3,4 and 6 disease manifestations respectively (Figure S8B). The cases where multiple mutations lead to different disease manifestations were mostly associated with syndromes (i.e. diseases having multiple phenotypic expressions) or leading to diseases affecting the same organ/system. For residue positions, where multiple mutations lead to the same disease we examined possible causes. About 78% of these residues are structural neighbours (within 5 Å) with mutations leading to the same disease.

We checked if any of the mutations corresponded to gain of function (GOF)/loss of function (LOF) from GOF/LOF database [4] based on Human Gene Mutation database [5] and MAVEDB [6]. Only 157 mutations out of 1389 (in well modelled regions) were annotated as GOF/LOF (143 LOF and 14 GOF). 320 domains out of 1194 domains had multiple mutations in the same domain, of which 85 domains had these residue mutations clustered together (mutations within 5 Å of each other), suggesting that they may be gain of function mutations [7–9]. Mutations in these domains lead to various neurological, sensory, kidney, developmental problems etc. Of these 320 domains with multiple mutations, 4 domains had 13 (out of 14) GOF mutations present in the dataset. The rest of the domains either were LOF (22 mutations were assigned LOF) or were not assigned to either GOF/ LOF mutation in the database.

The different disease ontologies in our database were mapped to the Human Disease Ontology database [10] by mapping the MIM (Mendelian Inheritance in Man) terms [11]. When grouped based on anatomical entities, our dataset has maximum representation for nervous system disorders followed by musculoskeletal system disorders. We should bear in mind that the dataset is biassed as we are only examining proteins without structural homologues in the PDB.

**Text S4 - Reasons for choosing tools/algorithms/cutoffs**

1. 5 Å cutoff for choosing neighbours - We chose a 5 Å cutoff as this is the approximate minimum distance between two amino acid side chains [12] and we wanted to ensure stringency in our analysis. Furthermore distance cut offs between 4.5-5 Å had been previously used [13,14] to create residue contact networks to study the effect of mutations and prediction of functional sites etc .
2. ddG>1 for FoldX predicted destabilising mutations - A ddG value greater than 0 is an indication that the mutation could have an impact on stability. However, because of the inherent difficulties in calculating ddG values computationally we have set a more stringent cut off of ddG>1 to predict likely impact.
3. P2Rank for ligand binding site - P2Rank is one of the best performing tools for ligand binding site prediction and is used by PDBeKB as a resource for binding site annotations [15] and has been widely used by the community to predict ligand binding sites [16–18].
4. meta-PPISP for protein-protein interfaces - While a large number of individual predictors complement each other meta-predictors like meta-PPISP which combine the results of multiple techniques (cons-PPISP, PINUP, and ProMate) provide more confidence in results compared to predictions based on individual techniques [19] and meta-PPISP has been used to predict interface residues in recent studies [20,21].
5. Cell list algorithm - The cell list algorithm is used to find all atom pairs within a certain cut-off distance and has been widely used in molecular dynamics simulations and other tools to calculate atoms within a particular distance cut-off [22–25].
6. MutPred2- MutPred2 was chosen because it is one of the most recent pathogenicity predictors and could be run as a downloadable on the protein sequence dataset. Other tools needed information on the mutations at the genome level or did not have a version to run a custom mutation list.

**Text S5 - Characterization of disease mutations in various PANTHER class**

We inspected if there were differences in the 3 PANTHER protein classes that had at least 16 proteins and 30 mutations in their class (metabolite interconverting proteins, transporters, scaffold/adaptor proteins). The total number of disease associated mutations were 302, 165, 67 respectively. We did not consider other classes either because there were too few disease associated mutations or sequences. We calculated the percentage of disease associated residues for each PANTHER protein class that were predicted as conserved, ligand binding, interface, pathogenic, destabilising and buried in the AlphaFold models and compared them against all proteins in our dataset (Figure S17). All these properties were enriched in disease associated mutations in metabolite interconverting proteins as compared to all proteins (Figure S17). The metabolite interconverting proteins encompasses a large number of protein subclasses, which might lead to diverse functional requirements, leading to enrichment of all factors. We see a significant increase in disease associated mutations near ligand binding sites compared to all proteins (37% from 12%), because these proteins predominantly bind ligands to carry out their functions (Figure S17). The transporters had more disease associated mutations as buried, pathogenic or near predicted interfaces (Figure S17). The scaffold/adaptor proteins had higher percentage of residues as pathogenic, near conserved or predicted interfaces and a lower percentage of residues near ligand binding sites compared to all proteins (Figure S17). This could be because these proteins largely need to interact with other proteins to carry out their functions.

**Text S6 - Polymorphisms**

The number of residues with polymorphisms were 998. Of these only 539 positions were in protein domains with good quality. Of these, 109 were near conserved residues, 37 near predicted ligand binding site, 141 near predicted protein-protein interface. The total number of residue positions associated with polymorphism near a functional site was 219 out of 539. Polymorphisms in 141 residue positions were predicted as destabilising according to ddG calculations by foldX and 61 residues were buried. Only 82 residues were predicted as pathogenic according to MutPred2. We see that a larger percentage of disease associated mutations were closer to functional site, buried, destabilising and pathogenic as compared to polymorphisms.

We also checked the relative abundance of amino acids after polymorphism mutations (compared to SwissProt51 database) and we noticed that the enriched amino acids are mostly neutral like Cys, His, Met, Gln, Arg, Ser, The, Val, Trp (Figure S15). The amino acids that were enriched in disease associated mutations compared to SwissProt51 database are Cys, His, Pro, Arg, Ser and Trp.

**Text S7 - Humsavar**

We only used the disease type of variant for analysis of disease associated mutations. In addition, we also compared the disease variants with polymorphisms. The current version of humsavar mentions the mutations as LP/P (Likely pathogenic/pathogenic), LB/B (Likely benign/Benign) or US (uncertain significance). The LP/P variants correspond to disease variants in the database we used and LB/B correspond to polymorphisms.

**Text S8 - Predicted high ddG mutations**

The Beta-1,4 N-acetylgalactosaminyltransferase 1 (Q00973) protein has a mutation Arg300Cys which leads to a disease Spastic paraplegia 26, autosomal recessive (SPG26). Another protein Feline leukaemia virus subgroup C receptor-related protein 1 (Q9Y5Y0) mutation of Ala241Thr leads to the disease Posterior column ataxia with retinitis pigmentosa. This mutation does not lie near any predicted functional site, however it leads to a ddG change upon mutation of greater than 2.5 (as calculated using FoldX) which is highly destabilising (Figure S14).

**Text S9 - RosettaFold models could explain mutations but not AlphaFold**

There were a range of reasons for this difference between impacts reported by AlphaFold and RoseTTAFold models. In certain cases, small parts of the proteins had different orientations in the two models (Figure S13). In other cases the regions of the AlphaFold models either did not have a high pLDDT score for the residue/model or the mutated residue did not fall within the 5 Å of the predicted functional sites. In some cases, the sites in the AlphaFold models missed the prediction threshold for the functional site predictors. Slight changes in the RoseTTAFold models allowed these sites to pass the prediction threshold and hence the sites were predicted as functional sites in RoseTTAFold models but not the AlphaFold models.

**Text S10 - Modelling across different superfamilies**

We wanted to check if certain superfamilies were modelled better than others. For this purpose, we used all the models available from the AlphaFold Protein Structure Database, we calculated the mean model quality for each of the CATH superfamilies. Only 3.5% of the superfamilies had a mean model quality score less than 70, indicating most superfamilies were modelled well irrespective of their type (Figure S2).

**Text S11 - Model quality vs Length**

The AlphaFold model quality did not depend on the length of the domain (correlation coefficient between length and model quality of 0.16, -0.13, -0.03 for CATH, Pfam and unassigned domains) (Figure 7). However, the model quality was lower for longer Pfam and unassigned RoseTTAFold domain models (correlation coefficient between length and model quality of 0, -0.35, -0.52 for CATH, Pfam and unassigned domains respectively) (Figure S6).

**Text S12 - Assignment of unassigned domains to cath**

With improvements in deep learning techniques and sequence embeddings, we are developing tools to classify very remote homologues into CATH [26]. Preliminary results show that 41 of the unassigned domains could be assigned to CATH superfamilies using the CATH HMM and structure comparison protocols [27]. For the majority of the remaining sequences more powerful embedding strategies [28] suggest a match to a possible superfamily but detailed manual analysis is underway to validate the assignments. This classification process will be accelerated once all the good quality AlphaFold models are brought in over the next year as these will help to validate the very remote homologues. 11 domains could be new folds but until we have brought all the AlphaFold structures into CATH, it is difficult to determine whether the unclassified domains are new folds or very remote homologues.

**Text S13 - Definition of terms**

1. FunFam - FunFams are coherent subsets of sequences predicted to have similar functions according to conserved specificity-determining positions in their multiple sequence alignment.
2. lDDT - The Local Distance Difference Test (lDDT) [29] is a superposition-free score that evaluates local distance differences of all atoms in a model. This score provides the per residue confidence in the prediction and is provided by both AlphaFold and RoseTTAFold as a measure of accuracy of the residue prediction. lDDT is also used by CAPRI to score protein models[30]

3. Neff - The number of effective sequences (Neff) is defined as the number of sequence clusters in the multiple sequence alignment after removing the rows and columns with greater than 25% gaps and clustering the remaining sequences using an 80% sequence identity cutoff.
4. Scorecons - Scorecons [31] is an entropy-based method that calculates the conservation of each position in the alignment with 0 being completely unconserved and 1 being completely conserved.
5. Percent scorecons - Percent scorecons is defined as the percentage of positions (calculated with respect to mean length of the sequences in the alignment) that have a scorecons value >=0.8. This measure provides an estimate of the percentage of conserved positions in the alignment.
6. DOPs - The DOPs (Diversity of Positions) score calculates the diversity based on the conservation scores (scorecons) and their frequencies. DOPs provides an overall conservation score for the entire alignment with a score of 0 indicating no diversity (all positions having the same scorecons value) and 100 indicating no positions having the same scorecons value i.e highly diverse.

**Text S14 - Methods**

1. MSA generation of protein domains - The seed alignments were enriched with homologs by performing iterative sequence *hhblits* [32] searches against uniclust30 [33] and BFD [34,35] databases. In the course of iterations, we generated four alignments corresponding to e-value cutoffs 1e-30, 1e-10, 1e-6 and 1e-3. The seed alignment from MAFFT was used as the input to the first iteration; each subsequent iteration was initialised with the alignment obtained from the previous iteration followed by filtering at 90% sequence identity and 50% coverage cutoffs. For the final alignment, we selected the alignment with the lowest e-value which met one of the two criteria: at least 2,000 sequences with 75% coverage or 5,000 sequences with 50% coverage (both at 90% sequence identity cutoff) were collected.
2. MSA for prediction of conserved sites - FunFams have been shown to produce MSAs that can be used to identify conserved residues highly enriched in functional sites [36] Sometimes these alignments are not large enough to be informative enough to detect conserved residues. Wherever possible we created MSAs (such that Neff>30, DOPs>70 and percent scorecons<20%) by either combining the closely related functional families or aligning the query sequence with members within its superfamily. The FunFam of the queried domain was merged to other FunFams in the superfamily (provided that the average length is within 80% of the queried FunFam), provided the e-value cut off of 1e-03 was met and the Neff of the alignment increased. This process was iterated until Neff was maximised. The domains whose MSA could not be expanded by merging FunFams were searched using HMMER3 within the specific superfamily with an e-value cut off of 1e-03. These alignments were used if the DOPs>70, percent scorecons<20% and Neff>30. For the remaining domains, the conserved positions were identified using the MSA used to build the models, provided they followed the DOPs and percent scorecons cut-off. These MSAs were built using metagenomic sequences and therefore tended to be much larger which sometimes might affect the detection of conserved sites.

### References


1. Mi H, Muruganujan A, Casagrande JT, et al. Large-scale gene function analysis with the PANTHER classification system. Nat. Protoc. 2013; 8:1551–1566
2. Jumper J, Evans R, Pritzel A, et al. Highly accurate protein structure prediction with AlphaFold. Nature 2021; 596:583–589
3. Baek M, DiMaio F, Anishchenko I, et al. Accurate prediction of protein structures and interactions using a three-track neural network. Science 2021; 373:871–876
4. Sevim Bayrak C, Stein D, Jain A, et al. Identification of discriminative gene-level and protein-level features associated with pathogenic gain-of-function and loss-of-function variants. Am. J. Hum. Genet. 2021; 108:2301–2318
5. Stenson PD, Mort M, Ball EV, et al. The Human Gene Mutation Database (HGMD®): optimizing its use in a clinical diagnostic or research setting. Hum. Genet. 2020; 139:1197–1207
6. Esposito D, Weile J, Shendure J, et al. MaveDB: an open-source platform to distribute and interpret data from multiplexed assays of variant effect. Genome Biol. 2019; 20:223
7. Campbell AJ, Watts KJ, Johnson MS, et al. Gain-of-function mutations cluster in distinct regions associated with the signalling pathway in the PAS domain of the aerotaxis receptor, Aer: Signalling in the Aer-PAS domain. Mol. Microbiol. 2010; 77:575–586
8. Kamburov A, Lawrence MS, Polak P, et al. Comprehensive assessment of cancer missense mutation clustering in protein structures. Proc. Natl. Acad. Sci. 2015; 112:E5486–E5495
9. Meyer MJ, Lapcevic R, Romero AE, et al. mutation3D: Cancer Gene Prediction Through Atomic Clustering of Coding Variants in the Structural Proteome. Hum. Mutat. 2016; 37:447–456
10. Schriml LM, Munro JB, Schor M, et al. The Human Disease Ontology 2022 update. Nucleic Acids Res. 2022; 50:D1255–D1261
11. . OMIM - Online Mendelian Inheritance in Man.
12. Jack BR, Meyer AG, Echave J, et al. Functional Sites Induce Long-Range Evolutionary Constraints in Enzymes. PLOS Biol. 2016; 14:e1002452
13. Prabantu VM, Naveenkumar N, Srinivasan N. Influence of Disease-Causing Mutations on Protein Structural Networks. Front. Mol. Biosci. 2021; 7:620554
14. Chakrabarty B, Parekh N. NAPS: Network Analysis of Protein Structures. Nucleic Acids Res. 2016; 44:W375–W382
15. PDBe-KB consortium, Varadi M, Berrisford J, et al. PDBe-KB: a community-driven resource for structural and functional annotations. Nucleic Acids Res. 2020; 48:D344–D353
16. Tunyasuvunakool K, Adler J, Wu Z, et al. Highly accurate protein structure prediction for the human proteome. Nature 2021; 596:590–596
17. Yang J, Kwon S, Bae S-H, et al. GalaxySagittarius: Structure- and Similarity-Based Prediction of Protein Targets for Druglike Compounds. J. Chem. Inf. Model. 2020; 60:3246–3254
18. Singh N, Decroly E, Khatib A-M, et al. Structure-based drug repositioning over the human TMPRSS2 protease domain: search for chemical probes able to repress SARS-CoV-2 Spike protein cleavages. Eur. J. Pharm. Sci. 2020; 153:105495
19. Xue LC, Dobbs D, Bonvin AMJJ, et al. Computational prediction of protein interfaces: A review of data driven methods. FEBS Lett. 2015; 589:3516–3526
20. Lo Gullo G, De Santis ML, Paiardini A, et al. The Archaeal Elongation Factor EF-2 Induces the Release of aIF6 From 50S Ribosomal Subunit. Front. Microbiol. 2021; 12:631297
21. Diesterbeck US, Gittis AG, Garboczi DN, et al. The 2.1 Å structure of protein F9 and its comparison to L1, two components of the conserved poxvirus entry-fusion complex. Sci. Rep. 2018; 8:16807
22. Soni N. neeleshsoni21/Cell_list. 2021;
23. Yao Z, Wang J-S, Liu G-R, et al. Improved neighbor list algorithm in molecular simulations using cell decomposition and data sorting method. Comput. Phys. Commun.

2004; 161:27–35
24. Dobson M, Fox I, Saracino A. Cell List Algorithms for Nonequilibrium Molecular Dynamics. ArXiv14123784 Phys. 2014;
25. Dhawanjewar AS, Roy AA, Madhusudhan MS. A knowledge-based scoring function to assess quaternary associations of proteins. Bioinformatics 2020; 36:3739–3748
26. Nallapareddy V, Bordin N, Sillitoe I, et al. CATHe: Detection of remote homologues for CATH superfamilies using embeddings from protein language models. 2022;
27. Sillitoe I, Bordin N, Dawson N, et al. CATH: increased structural coverage of functional space. Nucleic Acids Res. 2021; 49:D266–D273
28. Elnaggar A, Heinzinger M, Dallago C, et al. ProtTrans: Towards Cracking the Language of Life's Code Through Self-Supervised Deep Learning and High Performance Computing. 2020;
29. Mariani V, Biasini M, Barbato A, et al. lDDT: a local superposition-free score for comparing protein structures and models using distance difference tests. Bioinformatics 2013; 29:2722–2728
30. Kwon S, Won J, Kryshtafovych A, et al. Assessment of protein model structure accuracy estimation in CASP14 : Old and new challenges. Proteins Struct. Funct. Bioinforma. 2021; 89:1940–1948
31. Valdar WSJ. Scoring residue conservation. Proteins Struct. Funct. Genet. 2002; 48:227–241
32. Steinegger M, Meier M, Mirdita M, et al. HH-suite3 for fast remote homology detection and deep protein annotation. BMC Bioinformatics 2019; 20:473
33. Mirdita M, von den Driesch L, Galiez C, et al. Uniclust databases of clustered and deeply annotated protein sequences and alignments. Nucleic Acids Res. 2017; 45:D170–D176
34. Steinegger M, Mirdita M, Söding J. Protein-level assembly increases protein sequence recovery from metagenomic samples manyfold. Nat. Methods 2019; 16:603–606
35. Steinegger M, Söding J. Clustering huge protein sequence sets in linear time. Nat. Commun. 2018; 9:2542
36. Das S, Lee D, Sillitoe I, et al. Functional classification of CATH superfamilies: a domain-based approach for protein function annotation. Bioinformatics 2015; 31:3460–3467
